# Metacognitive Computations for Information Search: Confidence in Control

**DOI:** 10.1101/2021.03.01.433342

**Authors:** Lion Schulz, Stephen M. Fleming, Peter Dayan

**Affiliations:** Department of Computational Neuroscience, Max Planck Institute for Biological Cybernetics, University College London; Department of Experimental Psychology, University College London; Wellcome Centre For Human Neuroimaging, University College London; Max Planck Centre For Computational Psychiatry and Ageing Research, University College London; Eberhard Karls University of Tübingen

**Author notes:** Correspondence concerning this article should be addressed to: Lion Schulz, Department of Computational Neuroscience, Max Planck Institute for Biological Cybernetics, Max-Planck-Ring 8, 72076 Tübingen, Germany.

**Keywords:** Computation, confidence, metacognition, information search, decision making

## Abstract

The metacognitive sense of confidence can play a critical role in regulating decisionmaking. In particular, a lack of confidence can justify the explicit, potentially costly, instrumental acquisition of extra information that might resolve the underlying uncertainty. Recent work has suggested a statistically sophisticated tapestry behind the information governing both the making and monitoring of choices. Here, we extend this tapestry to reveal extra richness in the use of confidence for controlling information seeking. We thereby highlight how different models of metacognition can generate diverse relationships between action, confidence, and information search. More broadly, our work shows how crucial it can be to treat metacognitive monitoring and control together.

---

After carefully deliberating between Scotland and the Cote d’Azur, you have decided to spend your next summer holiday in the north of Britain. You are rather confident about this choice. Before it comes to booking your train tickets, an article pops up in your feed: “Skye or Saint-Tropez – the ultimate comparison”. Do you spend money and time on reading this article? Or do you purchase the tickets right away? Conflicts like this are sadly commonplace: Do you read another news story before heading to the polls? Do you consult a doctor before heading to the pharmacy? Regardless of the specific situation, we have to balance accuracy with (monetary or temporal) cost (Cohen, McClure, & Yu, 2007; Dayan & Daw, 2008). The decision to gather further information rather than making the choice based on our current knowledge thus depends critically on the initial choice’s expected rectitude, given current information – which is a form of subjective confidence (Pouget, Drugowitsch, & Kepecs, 2016).

Computational and cognitive neuroscience has extensively studied many aspects of such decisions (Gottlieb & Oudeyer, 2018; Gottlieb, Oudeyer, Lopes, & Baranes, 2013; E. Schulz & Gershman, 2019). For instance, a coupling between expected accuracy and information search is implicit in drift-diffusion models. There, participants can infer their chance of being correct from information that accumulates over time, and have to decide whether to stop or continue sampling evidence (Gold & Shadlen, 2001, 2007; Ratcliff & Rouder, 1998; Wald, 1949). These models have been highly successful in explaining the latent speed-accuracy trade-off present in many perceptual tasks (Bogacz, Wagenmakers, Forstmann, & Nieuwenhuis, 2010; Ratcliff, Smith, Brown, & McKoon, 2016).

However, humans also enjoy a more sophisticated and explicit sense of their expected accuracy, in forms of metacognition (Fleming & Daw, 2017; Shekhar & Rahnev, 2020; Yeung & Summerfield, 2012). Explicit metacognition refers to conscious representations of performance that are available for flexible usage in behavioural control or communication to others (Shea et al., 2014). In turn, these representations can be coupled to richer, normative, approaches to information search. In this paper, we provide a unified theoretical treatment of this relationship.

Metacognitive evaluations accompany a wide range of decisions, from basic judgments of perception and memory to reflective evaluations of our knowledge or the “goodness” of subjective choices (De Martino, Fleming, Garrett, & Dolan, 2013; Fischer, Amelung, & Said, 2019; Nelson & Narens, 1990; Rahnev et al., 2020). In turn, recent research has begun to reveal constraints on how metacognitive judgments are formed – both in terms of within-subject decision processes and between-subject factors (Fleming & Daw, 2017; Shekhar & Rahnev, 2020; Yeung & Summerfield, 2012). For example, in perceptual decision-making, human confidence judgments are influenced both by the uncertainty of sensory information, and the difficulty of making a particular discrete choice (Bang & Fleming, 2018; Pouget et al., 2016). Of particular interest are the processes that contribute to drops in confidence following errors. Such error monitoring can occur even in the absence of external feedback, and can rely on a purely internal evaluation mechanism (Boldt & Yeung, 2015; Rabbitt, 1966; Yeung, Botvinick, & Cohen, 2004). Moreover, confidence judgements also differ substantially between individuals, indicating personal-level influences on metacognition – a finding with implications for phenomena ranging from psychiatric disorders to political radicalisation (David, Bedford, Wiffen, & Gilleen, 2012; Hoven et al., 2019; Rollwage et al., 2018).

Theories of confidence have duly attempted to address these diverse aspects. Suggestions range from Bayesian accounts in which the same information underlies both decision and confidence (Cartwright & Festinger, 1943; Kepecs, Uchida, Zariwala, & Mainen, 2008; Sanders, Hangya, & Kepecs, 2016) to the possibility that extra inputs are available for the confidence rating that, for instance, accrue after or in parallel to a decision being made (Moran, Teodorescu, & Usher, 2015; Navajas, Bahrami, & Latham, 2016; Pleskac & Busemeyer, 2010). Broader notions of covariance between the information underlying both decision and confidence can capture both these aspects (Fleming & Daw, 2017; Jang, Wallsten, & Huber, 2012).

Information-seeking behavior is similarly highly complex and differs substantially between individuals. This again has implications, for example for psychiatric symptoms such as paranoia (Ermakova et al., 2018; Garety & Freeman, 2013; So, Siu, Wong, Chan, & Garety, 2016), or for patients suffering from obsessive compulsive disorder (Baranski & Petrusic, 2001; Hauser, Moutoussis, Dayan, & Dolan, 2017; Navajas et al., 2016; Tolin, Abramowitz, Brigidi, & Foa, 2003). Inter-individual differences in information search are also linked to real-world attitudes, as is evident in a relationship between lowered search and dogmatism (L. Schulz, Rollwage, Dolan, & Fleming, 2020).

Critically, metacognitive judgements are known to exert a causal influence over the choice to collect more information. For instance, in a study of perceptual decision-making, Desender, Boldt, and Yeung (2018) used a perceptual manipulation to induce different levels of confidence in different conditions, while keeping subjects’ objective performance equal. In the condition with lower confidence, subjects were more likely to seek additional information, providing key causal evidence for the role of confidence in the collection of information. In the memory domain, artificially boosting people’s confidence when learning word pairs makes them less likely to choose to study those pairs again, even though performance remains unchanged (Metcalfe & Finn, 2008). Other studies also support this close relationship between confidence and information search. For example, neural markers of confidence have been linked to variability in information search (Desender, Murphy, Boldt, Verguts, & Yeung, 2019) and different forms of confidence are proposed to influence the trade off between exploring new options and exploiting old ones (Boldt, Blundell, & De Martino, 2019; Wilson, Geana, White, Ludvig, & Cohen, 2014; Wu, Schulz, Speekenbrink, Nelson, & Meder, 2018).

While a rich literature exists modelling the processes supporting explicit confidence formation on the one hand and information search on the other, these two phenomena have yet to be related within a unified framework. Given the close coupling between metacognitive monitoring and control (Nelson & Narens, 1990), we here examine the implications of different theories of confidence for how subjects should elect to collect more information. We start by introducing the core components of information seeking at an abstract level, outlining the intuitions behind the relationships between action, confidence and information search. We then zoom in on these computations in more detail, first discussing different theories of monitoring as delineated by Fleming and Daw (2017) before considering what kind of downstream consequences arise in optimal control computations for information seeking. Finally, we investigate the resulting behavioral predictions.

## The Information-Seeking Problem

### General overview

Action, confidence, and information seeking can be investigated in minimal settings such as the bare-bones perceptual task presented in Figure 1A. There, participants are presented with a noisy stimulus (two boxes each with a different number of flickering dots), about which they have to first make an initial binary decision (more dots in the left or right box). They then express their confidence in this decision. Following this, they can decide whether to (1) see another helpful stimulus before making a final decision or whether they want to (2) make this final decision without any additional evidence. Seeing the second sample is associated with a cost, and the final (and possibly the initial) decision is rewarded.

**Figure 1.**
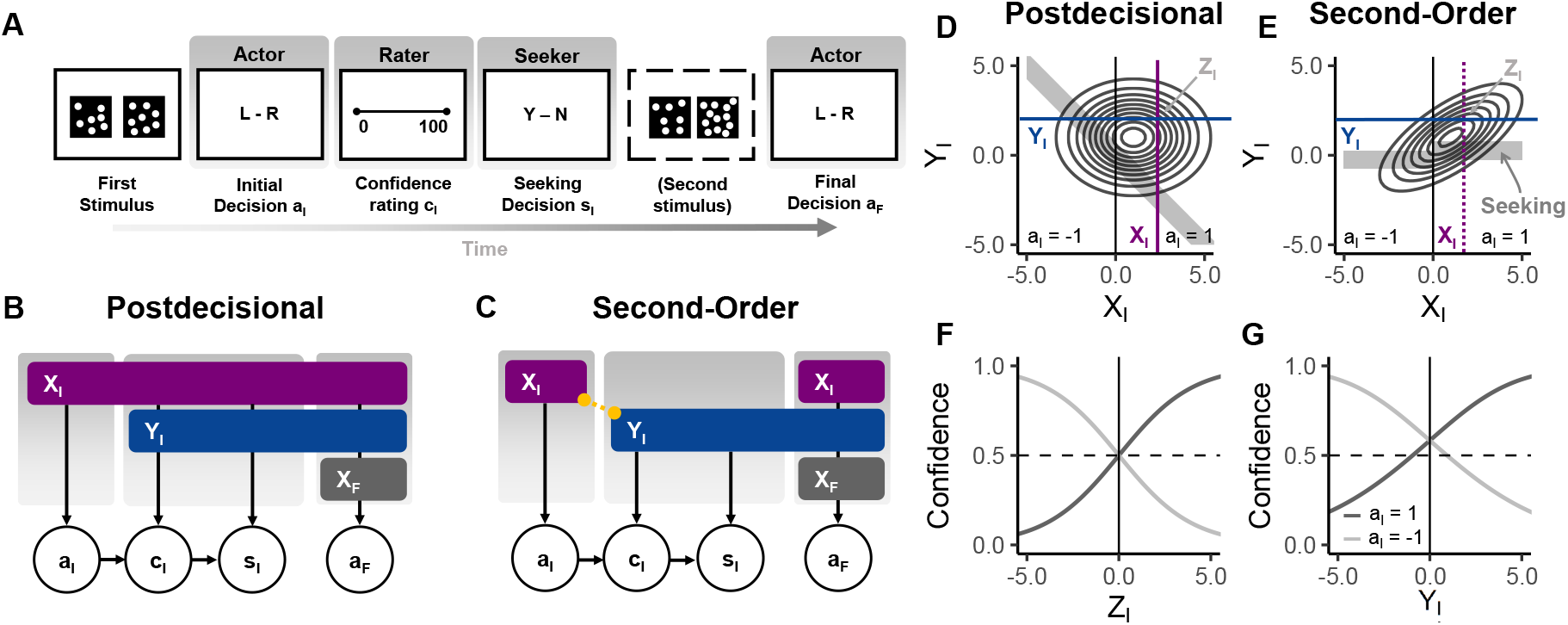
Example task and schematic of information available for different actions. (A) A task encapsulating the information-seeking problem presents a subject with a binary discrimination stimulus (more dots on the left, d = −1, or right, d = 1) about which it has to take and report a decision a_I_ and express its confidence in this decision c_I_. It can then decide whether or not (s_I_) it wants to see another additional stimulus which it could then use to make its final decision a_F_. We can conceptualize these sub-tasks as being made by three different agents with differing information. The actor makes a_I_ and a_F_, the rater expresses c_I_ and the seeker decides s_I_. (B,C) The information available for these different actions varies between the models: (B) In the postdecisional model, the actor takes a_I_ based on X_I_ and the rater and seeker have access to this X_I_ and an additional cue Y_I_ to rate the confidence and make the seeking decision. If the seeker decides to seek, the actor additionally receives X_F_ for the final decision. (C) In the second order model, the rater and seeker merely have access to Y_I_ (which can be correlated with X_I_) and a_I_ for c_I_ and s_I_. Across models, the final decision is made based on X_I_, Y_I_ and, depending on the seeking, X_F_. (D-E) Example stimulus distributions for d = 1 with example values for X_I_ and Y_I_ (and their sufficient statistic for d, namely Z_I_) highlighted. Within the distributions, we highlight zones in which the seeker would decide to seek out further information in grey. In the postdecisional model, this is a function of X_I_ and Y_I_, and in the second-order model a function of Y_I_, and a_I_. (F-G) Example postdecisional (F) and second-order confidences as a function of the relevant cue. Both support error monitoring by allowing confidence to be lower than 50 %. Note how in the second-order model the action has a boosting influence, for example increasing confidence above 0.5 for entirely ambiguous values of Y_I_.

Such a set-up is similar to the controlled environments previously used to study confidence and information seeking (Desender et al., 2018; Desender, Murphy, et al., 2019; L. Schulz et al., 2020). In these tasks, human subjects have been shown to modulate their seeking decisions based on their confidence, and also be sensitive to the cost of the additional information.

For illustrative purposes, we assign the paradigm’s different subtasks to three notional agents. These agents have, depending on the underlying confidence model, access to different information. The *actor* makes the two “objective” decisions (left or right). The *rater* expresses its confidence in these decisions (for brevity we only consider a first confidence rating here). A final agent, the *seeker*, is responsible for deciding whether additional information should be sought out (to improve the final decision of the *actor*). This terminology adds the seeker to the description of Fleming and Daw (2017), and makes it straightforward to specify the information that is available at each point in time and for each computation throughout the task. As we shall see, working with a concrete task forces a set of choices, for example, that the second choice of the actor can be informed by the confidence report of the rater. These will turn out to have a substantial impact on the results (for instance, that the more accurate the rater, the less information seeking is required).

In these terms: first, the rater perceives some evidence *X_I_*, and then makes a decision, *a_I_* ∈ {−1, +1}. The rater then publicly expresses its confidence, *c_I_* ∈ [0,1] in this decision, based on the information to which it has access. This information may or may not include *X_I_* and/or some unique information of its own *Y_I_*. Third, the seeker decides whether more information should be sought (*s_I_* ∈ {0,1}). The actor then makes a final decision, *a_F_* ∈ {−1, +1}. In our simple formulation, *a_F_* can be based on *X_I_* along with *c_I_* (since the rater’s confidence judgment is veridical and public) and, if extra information was sought, a further sample, *X_F_*. We refer the reader to Table 1 for an overview of the notation used throughout.

**Table 1.** Notation. We distinguish between (A) aspects of the task, (B) random variables, and (C) quantities/actions that the agent computes

| Notation | Explanation |
| --- | --- |
| **A. Experiment** |  |
| **States** |  |
| $I_d \in \{I_{-1}, I_1\}$ | Initial actual state of the problem |
| $F_d \in \{F_{-1}, F_1\}$ | Final state of the problem |
| $d \in \{-1, 1\}$ | Underlying state of the stimulus |
| **Rewards and costs** |  |
| $r_I, r_F$ | Reward for the initial and final decisions |
| $r_S$ | Cost for obtaining the additional stimulus $X_F$ |
| **B. Random variables and their attributes** |  |
| **Random variables** |  |
| $X_I, X_F$ | Actor's stimuli at $I_d$ and $F_d$ |
| $Y_I$ | Rater's stimulus at $I_d$ (in postdecisional and second-order model) |
| $Z_I$ | Combination of $X_I$ and $Y_I$ |
| $Z_F$ | Combination of $X_I, Y_I$ and $X_F$ |
| **Noise terms associated with random variables** |  |
| $\sigma_I, \sigma_F$ | Standard deviation of $X_I$ and $X_F$ |
| $\tau_I$ | Standard deviation of $Y_I$ |
| $\zeta_I, \zeta_F$ | Standard deviation of $Z_I$ and $Z_F$ |
| $\rho_I$ | Correlation between $X_I$ and $Y_I$ |
| $\Sigma_I$ | Covariance matrix for $X_I$ and $Y_I$ with $\sigma_I, \tau_I$ and $\rho_I$ |
| **C. Agent** |  |
| **Actions and expressions** |  |
| $a_I, a_F \in \{-1, 1\}$ | The actor's initial and final decisions |
| $c_I \in [0, 1]$ | The rater's confidence in the actor's initial decision |
| $s_I \in \{0, 1\}$ | The seeker's decision whether (1) or not (0) to seek |
| $a_{F, s_I}$ | The actor's final decisions conditioned on the seeking decision |
| **Values and action values** |  |
| $Q_S(s_I)$ | Action values for seeking $Q_S(1)$ or not seeking $Q_S(0)$ |
| $V_{F, Z_F}^*$ | Value of having a specific cue $Z_F$ at final state $F$ |
| $V_{F, Z_I, s_I}^*$ | Value of having a specific cue $Z_I$ at final state $F$, conditioned on whether agent will seek or not |
| $V_{F, Y_I, s_I}^*$ | Value of having a specific cue $Y_I$ at final state $F$, conditioned on whether agent will seek or not |

We now unpack the computations behind these steps further, first discussing the initial decision and confidence, as outlined by Fleming and Daw (2017), before elucidating their consequences for the seeker’s decision.

### Formalising action and metacognitive monitoring

We can frame the decision-making problem as a partially observable Markov decision problem, or POMDP (Monahan, 1982; Sutton & Barto, 2018). In this, the actor’s first task is to use its cue *X_I_* to infer which of two states of the world *I_d_* (with *d* ∈ {−1,1}) it inhabits. These two states can represent a multitude of stimuli and task configurations, including more dots in the left (*I*_−1_) or right (*I*_1_) box in the task of Figure 1A, but equally any other binary judgement. The actor’s cue *X_I_* only affords partial information about d and is conventionally thought to be drawn from a normal distribution with mean d and standard deviation *σ_I_*.

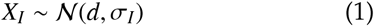

In a task only capturing the first decision, we might incentivize the initial decision *a_I_* with a pay-off of *r_I_* points for correct choices and 0 points for incorrect choices. In this case, the actor should optimally compare its sensory sample against a threshold. Under our stimulus and pay-off regime with equal noises and pay-offs and equally prevalent underlying states, this threshold is optimally set to 0, implying that:

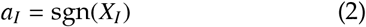

More complex schemes for pay-offs (e.g. more reward for correctly identifying *d* = 1) or asymmetric sources will impact this decision rule (Dayan & Daw, 2008), but we will focus on this simple set-up for clarity. In general, the expected performance of the actor is determined by *σ_I_*, with higher values associated with more mistakes, on average (see below for more details).

The rater’s task is now to compute a confidence, *c_I_*, in *a_I_*. We assume that the rater follows Bayesian precepts and reports its belief that *a_I_* was the correct choice, given its information (which we here denote by *C*) and the task parameters *θ*:

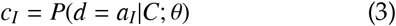

Fleming and Daw (2017) discuss three different models for the nature of *C*, the first-order, the post-decisional and the second-order model. We now recapitulate these models before adapting them to the information-seeking problem. Since the first-order model is a special case of the postdecisional model, we discuss them jointly.

#### Postdecisional and first-order models

In the *postdecisional* model, the rater knows the actor’s information *X_I_* and action *a_I_*. It also receives *independent* postdecisional information, *Y_I_* (see also Figure 1B). This postdecisional cue *Y_I_* is sampled from a distribution with the same mean d but with its own standard deviation *τ_I_*,

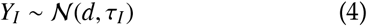

The first-order model is an instance of the postdecisional model in which *τ_I_* = ∞. In other words, in its case, the rater has no extra postdecisional information over and above the rater.

The rater first combines its sample with the actor’s sample in a precision weighted fashion, leading to a sufficient statistic *Z_I_* which has a standard deviation of *ζ_I_*:

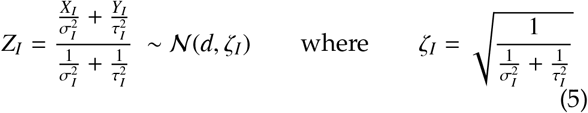

The rater’s confidence in the actor’s choice then comes from the posterior distribution obtained through Bayes’ rule. Here, the distance between the threshold and *Z_I_* becomes a proxy for the rater’s confidence:

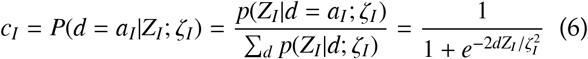

An important facet of the postdecisional model is that *Z_I_* and *a_I_* can “contradict” each other. In other words, the rater might have information that favours one judgement (e.g. *Z_I_* = 0.7) while the actor might have had information that favoured the other (e.g. *X_I_* = −0.2). Such a disagreement will lead the confidence to be lower than 0.5, triggering what is known as error monitoring (Boldt & Yeung, 2015; Fleming & Daw, 2017; Yeung & Summerfield, 2012) as we see in Figure 1F.

In the first-order model, with *τ_I_* = ∞, the actor and rater have the same information (*Z_I_* = *X_I_*), but the confidence computations outlined in equation 6 still hold. As a consequence, the rater will always endorse the actor’s choice in the first-order model. This, in turn, prevents it from exhibiting error monitoring. Furthermore, and inconsistent with empirical observations of dissociations between performance and metacognition (Rahnev et al., 2020; Shekhar & Rahnev, 2020), it ensures the actor and the rater’s accuracy remain coupled, as we will discuss in more detail below. ^1^

#### Second-order Model

In the postdecisional model, the rater is particularly well endowed with information: it knows exactly what the actor used to make its decision. This assumption might not hold under several assumptions, for example different neural pathways for action and confidence. Therefore, more general models allow merely for correlation between the information employed by actor and rater (Fleming & Daw, 2017; Jang et al., 2012). One example for such a set-up is Fleming and Daw’s (2017) *second-order model*. In it, the rater still receives *Y_I_*, but it is denied *X_I_*. Rather, it only observes the binary decision *a_I_* (see also Figure 1C). However, in a key contrast to the postdecisional model, the actor’s and rater’s information are correlated:

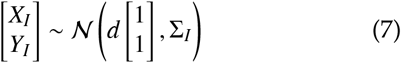

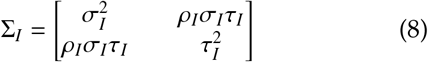

This correlation is visible in Figure 1E and allows the rater to make partial inferences about the location of *X_I_*. To compute its confidence, the rater thus combines two actual, and one inferential, source of information about *d*: the action *a_I_*, its own *Y_I_*, and the information provided by these variables about *X_I_* via the covariance between *X_I_* and *Y_I_*, to compute a posterior:

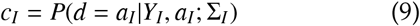

This expression involves more complex computations, including marginalization over possible values of *X_I_*, which we recapitulate and discuss in the appendix (section A). As with the postdecisional model, the second-order model also supports error monitoring, and can give rise to different levels of metacognitive insight.

A crucial aspect of the second-order model is that an agent (here, the rater) has to “[infer] the causes of its own action” (Fleming & Daw, 2017). This (partial) decoupling of action and confidence information gives it more flexibility than the postdecisional model and makes the action a crucial input to the computation, in turn boosting confidence for ambiguous *Y_I_*’s. This is visible in Figure 1G, and discussed at length in Fleming and Daw (2017)^2^.

### Formalizing the information-seeking problem

After a confidence estimate is formed, the seeker needs to decide whether the actor should see additional information before making its final decision *a_F_* about *d*. To conceptualize this more formally, we extend our POMDP (Dayan & Daw, 2008; Gottlieb et al., 2013) by adding a second pair of states *F_d_* that deterministically follow *I_d_* (*I*_−1_ → *F*_−1_, *I*_1_ → *F*_1_). If the seeker decides to seek, the actor receives a second stimulus *X_F_* at *F_d_* which it can use to make its final decision. We again assume this second cue to be sampled from a normal distribution with mean *d* and an associated standard deviation *σ_F_*:

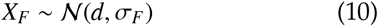

A final correct decision again comes with a remuneration *r_F_* whereas an incorrect choice leads to 0 points. Finally, and crucially, seeking incurs a cost, *r_s_* ^3^.

Regardless of the metacognitive information configurations outlined above, the seeker’s choice involves the same basic question: Is seeking worth the cost? To decide, it computes two action-values, *Q_S_* (*s_I_*): one for seeking *Q_S_* (1) and one for not seeking *Q_S_* (0). In short, these involve predicting how accurate the actor will be on the final decision, with or without *X_F_*. We will next outline these computations, first explaining the details in the simpler case of the postdecisional model, and then highlighting differences in second-order computations. Across these models, we assume that the seeker has the same information as the rater, i.e., representing different sides of the metacognitive coin (monitoring and control). This will allow us to capture key unique contributions of confidence to information search over and above the actor’s objective performance.

To illustrate the computations involved, we follow recent studies (Desender et al., 2018; L. Schulz et al., 2020) who provide no reward for the initial decision (*r_I_* = 0). We fix the reward for the final decision at *r_F_* = 1 and will show different costs for the additional stimulus *r_S_*. Furthermore, we assume a noisier first (*σ_I_* = 1.5) than second (*σ_F_* = 1) stimulus, similarly to L. Schulz et al. (2020).

#### Seeking in the postdecisional/first-order case

To compute the two *Q*-values, we first need to consider how the final decision *aF* is made at *Fd*, with and without *X_F_*.

If the seeker decides to collect no further information, the actor’s final decision will be based on the same information as the rater’s initial confidence. As a result, the final decision will just repeat its initial decision *a_F,0_* = *a_I_* if *Z_I_* and *X_I_* agree. If they contradict each other, the actor will correct what it assumes to be an initial mistake and change its mind. In confidence space, this transition occurs at *c_I_* = 0.5

To compute the associated action value for not seeking, *Q_I_*(0), the seeker first computes the optimal expected value 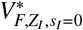 of having a specific *Z_I_* at *F_d_* conditioned on its non-seeking behavior. This involves multiplying the reward obtained through a correct final decision with the probability of making a final correct decision based on *Z_I_* (assuming that incorrect decisions incur no cost). In the postdecisional model, this probability is simply max{*c_I_*, 1 − *c_I_*}. Figure 2 shows the sub-components of the postdecisional seeker. For clarity, we there assume that *τ_I_* = ∞, reducing the problem to the first-order model. Figure 2A depicts the posteriors and the associated values. Importantly, this is the equivalent of the curves for the first-order confidence. Since *not* seeking costs nothing, the optimal action-value for not seeking is just this value:

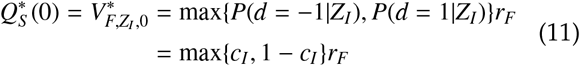

**Figure 2.**
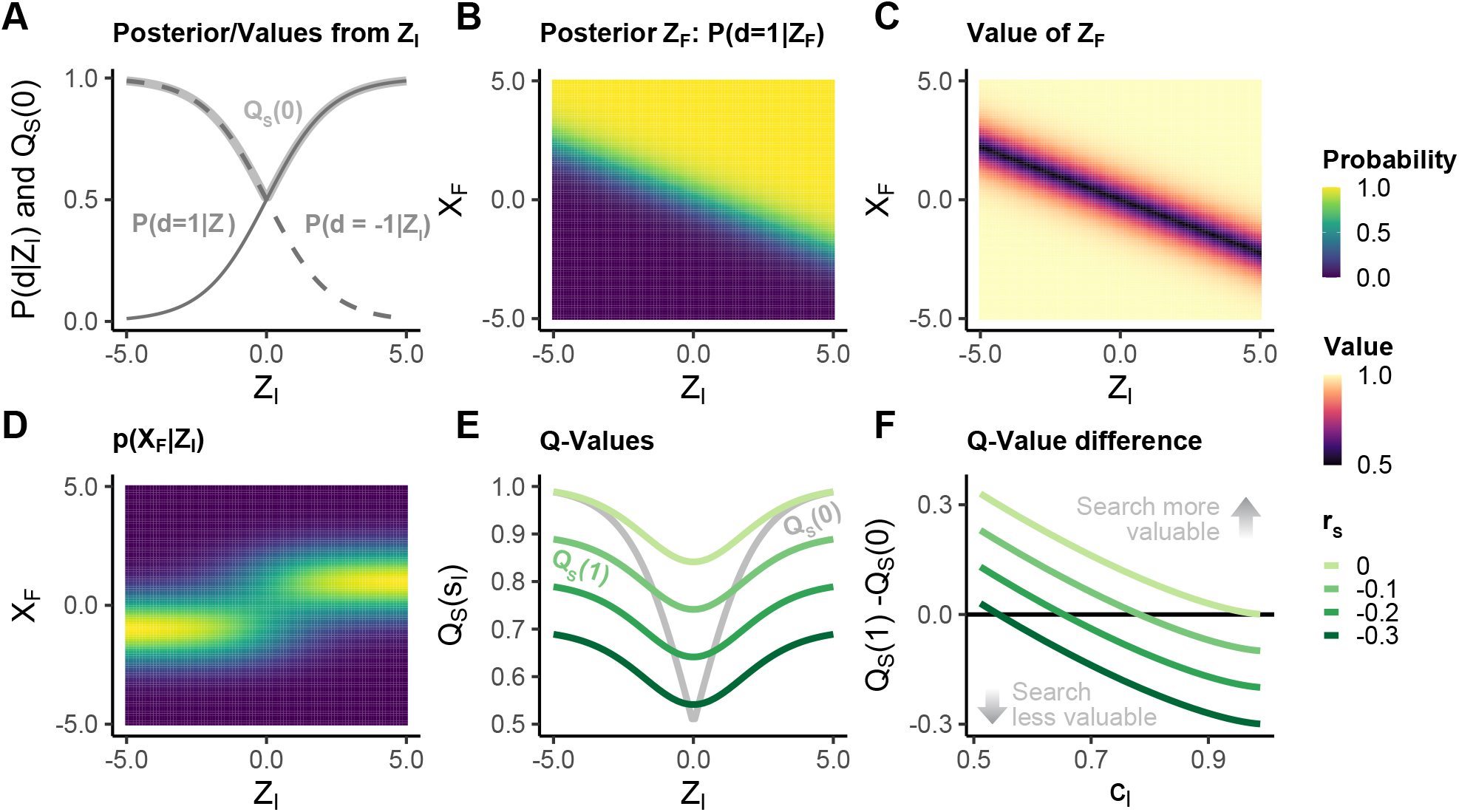
Subcomponents of the information-seeking computation in a first-order/postdecisional model. (A) To compute the Q-value of not seeking, Q_S_ (0) (bold line), the agent computes the max of the two posteriors over d from Z_I_, P(d = −1 |Z_I_) and P(d = 1 |Z_I_). (B) If the seeker decides to search, it receives X_F_ which it combines in a precision-weighted fashion with Z_I_ to form a Z_F_. Here, we plot the posterior P(d = 1|Z_F_). Because Z_I_ is noisier than X_F_ (ζ_I_ > σ_F_), the apparent slope of this posterior is not −1. Rather, X_F_ is weighted more than Z_I_. The converse posterior P(d = − 1|Z_I_) = 1 − P(d = 1|Z_I_) is the remaining probability. (C) The seeker computes the value associated with a given ZF from the maximum of these two possible posteriors. (D) Because it needs to decide whether to seek or not before receiving X_F_, the agent needs to predict X_F_. It does this by summing the two possible source distributions 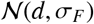 weighted by their individual confidence values. (E) To compute the value for seeking Q_S_ (0), the agent averages over the two quantities in C and D, based on its Z_I_. We here display the Q-value for not seeking and for seeking overlayed, with the latter shown as a function of the seeking cost r_S_. Note how the maximum of the Q-value for seeking Q_S_ (0) is defined by this cost. (F) The agent seeks when seeking is more valuable than not seeking. We here display the difference between the two values, transformed into confidence space. As confidence increases, the benefit of seeking decreases. Partially adapted from Dayan and Daw (2008). Parameters set at ζ_I_ = σ_I_ = 1.5, σ_F_ = 1.

In contrast, if the seeker decides to seek, the actor can use the additional stimulus *XF* to disambiguate *d* further for its final decision *a*_*F*,1_. It does this by first forming a final combined variable *Z_F_* in a precision weighted fashion equivalent to equation 5:

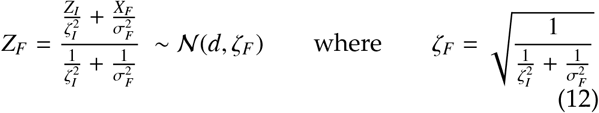

Similarly to the first decision, the actor can then compare *Z_F_* against a threshold (again, optimally *Z_F_* = 0 given our pay-off regime) to make the final decision. We plot the posterior associated with this value for *d* = −1 in Figure 2B. There, the threshold for *a_F_* lies on where *P*(*d* = 1 |*Z_F_*) = 0.5.

Here, the initial stimulus *Z_I_* is noisier than the final stimulus *X_F_* (σ_I_ = 1.5 and *σ_F_* = 1). As a result, *X_F_* is given more weight than *Z_I_* in the posterior. For example, an *X_F_* = 1 will increase the *P*(*d* = 1|*Z_F_*) posterior more than an equivalent *Z_I_* = 1. Similarly, a less extreme *X_F_* will be necessary to overturn a *Z_I_* of a different sign. This is evident in the tilt of the posterior, which is not fully diagonal but rather slants towards *X_F_*.

As for *Z_I_* in the no-seeking calculations, we compute the expected value of a given combination of *Z_I_* and *X_F_* from the maximum of the two possible posteriors (where *P*(*d* = − 1 |*Z_F_*) = 1 − *P*(*d* = 1 |*Z_F_*):

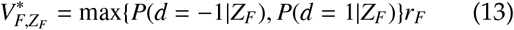

We plot this value 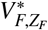 in Figure 2C as a function of *Z_I_* and *X_F_*. Again, the slope of the relationship is determined by the greater contribution to *Z_F_* of *X_F_* than that of *Z_I_*.

Crucially, however, the seeker has to decide whether it wants to seek *before* the actor has seen *X_F_*. It therefore needs to predict this second cue. The resulting distribution *p*(*X_F_*|*Z_I_*) is a function of how likely the seeker believes that the actor is to receive a stimulus from one of the two means, or a sum of the two possible source distributions weighted by the rater’s initial confidence *c_I_* (see appendix A). Figure 2D shows this distribution as function of *Z_I_*, a mixture of two Gaussians.

To compute the expected value, 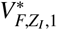, without having seen *X_F_*, the seeker then integrates over this distribution and the previously defined value function for its value of *Z_I_* given the prospect of seeking. Based on this mean value the seeker can now work out the action-value for seeking by considering the cost of the search:

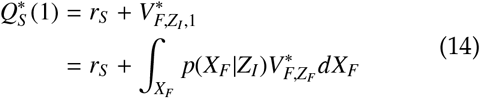

This value is shown in Figure 2E as a function of *Z_I_* and for different seeking costs *r_S_*. It is highest when the seeker expects the final choice to be likely correct, that is when it is relatively sure about the identity *X_F_*. With more ambiguous values of *Z__I__*, this prediction can only be made with less certainty. The ceiling of the seeking value is defined by the cost *r_S_*. We plot the value for not-seeking in Figure 2E. It approaches 0.5 as *Z_I_* becomes less distinctive, and *c_I_* therefore becomes lower. The larger of the two *Q*-values then determines the seeking choice:

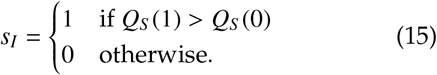

For ambiguous values of *Z_I_*, seeking is useful and will likely produce a better final outcome, even when taking into account the additional cost. When we transform the difference between the two values into confidence space (Figure 2F), we notice that, seeking will occur in lower confidence ranges, highlighting confidence’s crucial role in guiding the decision to seek.

#### Seeking in the second-order case

The second-order model entails some additional subtleties stemming from the different sources of information of actor, rater, and seeker. Recall that, in the second-order model, the rater only observes *a_I_* and *Y_I_* but does not have full access to the actor’s random variable *X__I__* (compare Figure 1C). Similarly, one might assume that the actor does not directly know *Y_I_* but only observes the rater’s utterance, *c__I__*. However, because the actor knows its own first action, *a_I_*, it can leverage the knowledge about the rater’s confidence algorithm to infer the initial confidence variable, *Y__I__*, underlying *c__I__*. It can then combine this random variable with *X_I_* to form *Z__I__*, taking into account the cues’ relative precisions and their covariance (see appendix A). The reason the actor can extract *Y_I_* from *c_I_*, but the rater cannot infer *X_I_* from *a_I_*. The confidence *c_I_* is continuous, whereas the action *a_I_* is discrete.

In the case of no seeking, the actor makes its decision based on *Z_I_* in a similar vein to the postdecisional model. In contrast to the postdecisional model, such a change of mind is however not necessarily coupled to *c__I__* < 0.5 given specific stimulus configurations, because of the additional information possessed by the actor at the second stage. Regardless, the value computations for holding a given *Z__I__* are equivalent to the post-decisional model (we detail this 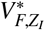 in the appendix section A). However, the seeker does not know *Z_I_*, because it does not have access to *X_I_*. It therefore has to marginalize out this quantity in a similar manner to the postdecisional model’s seeking computations:

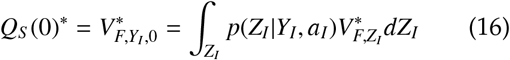

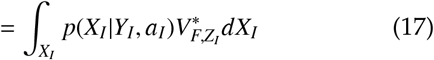

When the seeker decides to seek, the actor receives *X_F_* (again as per equation 10) which it combines with *Z_I_* to form a joint variable *Z_F_* (because there is no correlation between *X_F_* and *Z_F_* this is optimally done in a manner analogous to the postdecisional model, equation 12). This final variable *Z_F_* can then again be compared against a threshold for *a_F_*,_1_ and is used to compute a value. Similarly to the first-order and postdecisional models, the seeker does not know all the parts of *Z_F_* and has to marginalize over the unknowns. As before these mean values are then used to compute the *Q*-values associated with seeking and not seeking:

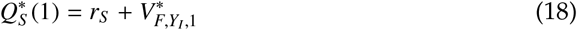

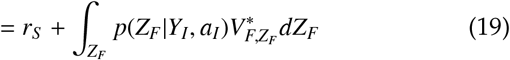

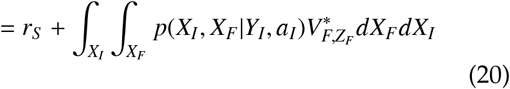

#### Commonalities and differences between the models’ seeking computations

While their details diverge, the different models still share some commonalities. Crucially, they all employ their current confidence to predict the future location of the second stimulus, *X_F_*, which is then combined with the final value of a stimulus combination to form the *Q*-value for seeking. For the postdecisional (and first-order) model confidence is also a determinant of the value of not seeking. The second-order model also uses the confidence to compute the non-seeking value, albeit by harnessing it to predict the location of *X_I_*, similarly to the postdecisional seeking value. All this highlights the crucial role of metacognition and confidence in the seeking decision.

## Results

In the following, we discuss features of these models in the information-seeking task. The task allows us to investigate several behavioral markers of action, confidence, and information search. With regard to the initial decision, we can observe (1) the average initial decision performance, (2) the initial confidence, and (3) an agent’s metacognitive accuracy (their ability to tell apart correct from incorrect choices through their confidence). With regards to the information-seeking decision, we can investigate both (1) the average level of information search as well as (2) the stopping criterion. Finally, we can observe how accurate an agent is. Our models predict specific patterns of interactions between these behavioural markers.

### Initial accuracy, average confidence and information seeking

Good information seeking should be coupled to decision accuracy: The more likely we are to make a mistake, the more additional information should help us. Broadly speaking, humans follow this prescription, seeking more information when they are worse at a task (Desender et al., 2018; Desender, Murphy, et al., 2019). Increased accuracy should also go hand in hand with a rise in average confidence, a corollary also supported by empirical evidence (Henmon, 1911; Nelson & Narens, 1990; Rahnev et al., 2020).

To investigate this in the context of our models, we fix the quality of the second stimulus as well as the cost for seeking, and investigate an agent’s average confidence and information search. We show these markers as a function of the initial accuracy which, across all models, is a function of *σ__I__*:

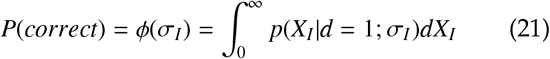

A figure showing this function is displayed in Figure 3A: The *lower* the actor’s noise *σ* becomes, the more accurate the objective decision.

**Figure 3.**
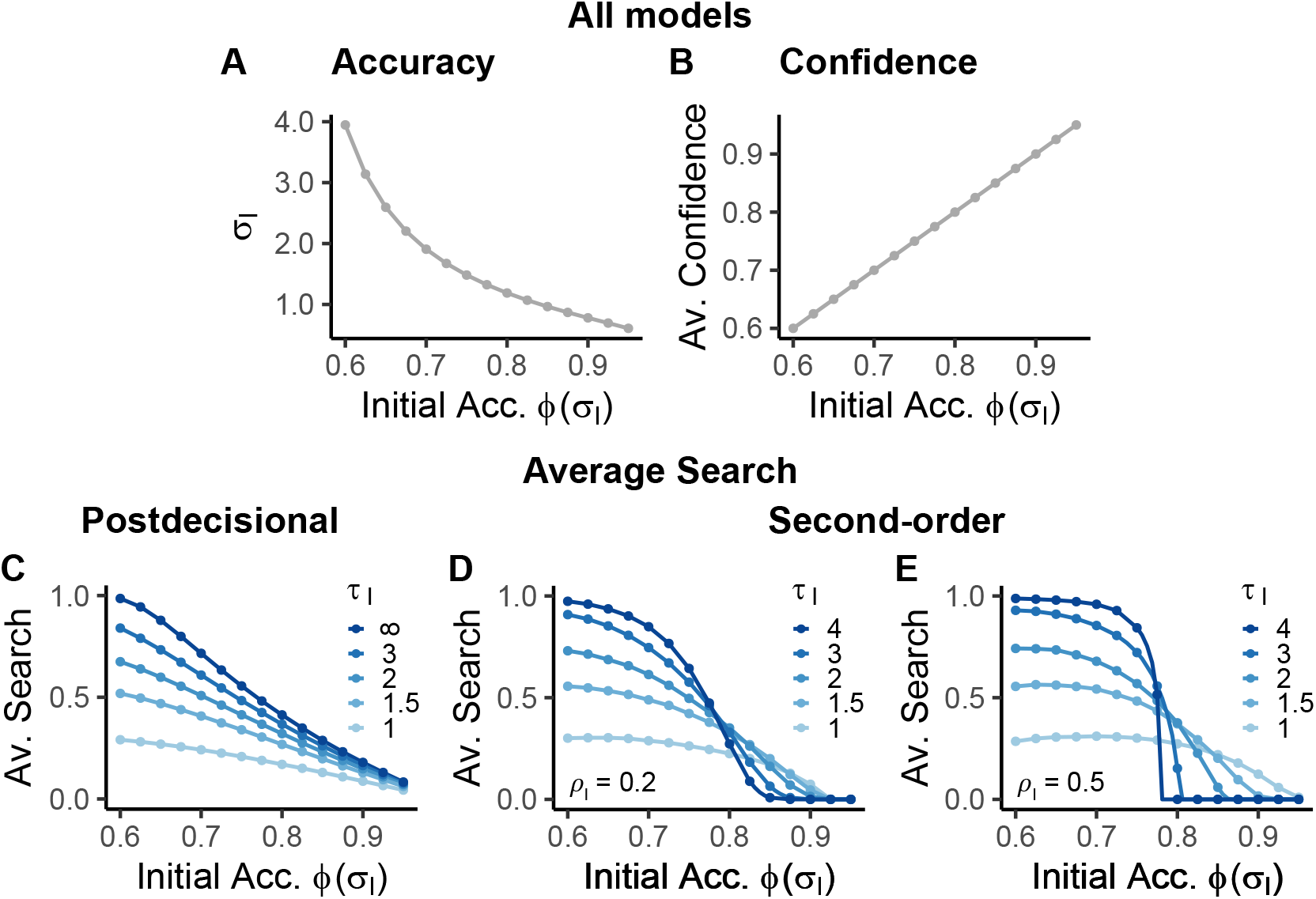
Initial accuracy, average confidence and information search. (A) Across models, the accuracy of the agent’s initial decision is governed by σ__I__ through the function ϕ(σ__I__) (B) In all models, the average initial confidence matches the average initial accuracy (C,D,E) Average seeking decreases with increasing initial accuracy for all models. However, the precision of the rater’s stimulus, τ__I__, moderates this relationship differently depending on the model. (C) In the postdecisional model, lower rater noise leads to less seeking across the accuracy spectrum because the rater and seeker have more additional information. (D, E) In the second-order model, high rater noise is associated with reduced seeking at higher levels of initial accuracy. This is because the seeker lacks direct access to the actor’s actual cue X__I__ and thus has to trust its decision. The correlation between the two (ρ__I__) modulates this effect (D: ρ__I__ = 0.2; E: ρ__I__ = 0.5). In the seeking plots, final stimulus noise and cost are fixed at σ_F_ = 1 (ϕ(σ_F_) = 0.84) and r_S_ = −0.1, respectively. The effect of ρ__I__ is shown further in section B of the appendix.

#### Average confidence across models

Across all our models, the average confidence is correctly calibrated and so tracks the objective accuracy (Figure 3B). For example, when the actor correctly responds in 71% of cases, the rater’s average confidence will also be 71%. As a result, simply measuring average confidence would allow no inference about the underlying model. Average confidence is also a coarse measure because it fails to describe the underlying confidence distribution well, as we will see further below.

#### Postdecisional and first-order model

Figure 3C shows how the first-order (τ*_I_* = ∞) and more general postdecisional models capture the relationship between accuracy and information seeking: the lower the initial accuracy, the more likely it is to seek out information. In fact, average search approaches an asymptote of 1 below a certain accuracy for the first-order model given the final stimulus precision and information cost used here. In other words, when objective accuracy is low it will almost always be worthwhile for a first-order agent to seek despite the cost.

The extra information provided by the postdecisional stimulus *Y_I_* impacts this average seeking over and above the average objective accuracy. Specifically, we see a marked reduction in seeking with lower *τ_I_* in comparison to the first-order model. This arises because the joint noise *ζ__I__* associated with *Z__I__* decreases as *Y__I__* becomes more precise, and since, in our model, *a_F_* can be informed by *Z_I_* even without extra seeking. In fact, the optimal cue-combination in equation 5 ensures that the joint noise ζ*_I_* will never be larger than the smallest of each of the two underlying variances; it will be smaller than the smaller of the two when the postdecisional noise is less than infinite. Consequently, a postdecisional seeker with *τ_I_* < ∞ always possesses information that is at least as accurate as a first-order seeker with an equivalent *σ_I_*. It will thus always seek out less information than its first-order sibling. The postdecisional noise *τ_I_* governs the difference between the pair with more precise postdecisional cues leading to larger differences.

This joint standard deviation *ζ_I_* also impacts the asymptote: specifically, the average propensity of a postdecisional agent to seek will never exceed a specific proportion (for *τ_I_* < ∞), even when its objective decision quality remains poor. This, and the generally lower search even before the asymptote, is partially due to the inherent capability for error monitoring in the postdecisional model: If the agent receives a postdecisional cue with sufficiently low noise that contradicts its initial cue, then it can infer that its initial choice was erroneous. When this postdecisional signal is strong enough to trigger a high error probability, then the agent can simply change its mind at *a_F_* without requiring the additional information. In contrast, when the actor has made an initially correct choice, a precise postdecisional stimulus will likely increase the rater’s confidence. The heightened confidence in turn will also decrease the need to seek additional information. Because the determinant of the average search is *ζ_I_*, and this quantity is close to *τ_I_* when *σ_I_* is large, a postdecisional agent’s average information seeking equals the average information seeking of a first-order agent whose actor noise *σ_I_* is equivalent to the postdecisional model’s rater noise *τ_I_*. In other words, when the actor *a_I_* knows almost nothing, the average seeking behaviour of a postdecisional agent will still resemble a first-order model whose decision accuracy would be governed by *τ_I_*.

#### Second-order model

The seeking behaviour of the second-order model differs in key aspects from the postdecisional model (see Figure 3D;E). While it also, broadly speaking, reduces its search with increasing initial accuracy, it exhibits a marked interaction between actor and rater noise. Specifically, the average seeking curves appear similar to those of the postdecisonal model when the initial accuracy of the second-order actor is relatively low. There, rater/seekers with higher *τ_I_* cues will seek more than those with lower *τ_I_*. Strikingly however, when the actor is more reliable (the initial accuracy is higher), second-order agents with higher *τ_I_* will seek less than those with more precise rater information.

The peculiar interaction between objective accuracy and confidence-noise arises from the second-order model’s informational set-up: Whereas the postdecisional model makes use of both *X_I_* and *Y_I_*, the second-order architecture only affords the rater access to a single cue, *Y_I_*, and the actor’s initial decision *a_I_*. This leaves it to make inevitably imperfect inferences about *X_I_*.

When the actor is relatively accurate (e.g. *σ_I_* = 1) and the rater’s information relatively inaccurate (e.g. *τ_I_* = 3), the rater has little information about the actor, but knows that the decision is likely correct (because *ϕ*(*σ_I_* = 1) = .84) – which will even be the case if the rater has an entirely ambiguous or even somewhat contradictory *Y_I_*. As a result, its confidence will remain high, even when *Y_I_* and *a_I_* contradict each other (see appendix B and Figure 6F). In other words, the rater will essentially resort to “trusting” the actor’s action across a wide range of its own information *Y_I_*. Because the seeker is equipped with the same information as the rater, it will likewise have too little information to justify the cost of seeking. Consequently, it will either fully trust or distrust the actor’s initial decision. In extreme cases, when *τ_I_* approaches ∞, the relation between initial accuracy and average seeking will in fact resemble a step-function.

Relatedly, there is a marked lack of seeking for high accuracies in the second-order model when keeping *τ_I_* constant. Notice how under the conditions of the cost of sampling and the accuracy of the second sample in Figure 3C, the first-order model will still search on up to a quarter of trials at 90% initial accuracy. In comparison, a second-order agent does not seek at all beyond that point with any but the most insightful values of *τ_I_* (Figure 3D;E). This again comes down to the fact that the rater and seeker have no alternative but to trust the actor’s decision when *τ_I_* ≫ *σ_I_*. When the objective accuracy is very high (e.g. *ϕ*(*σ_I_* = .8) ≈ 90%) such an imbalance arises even when the rater noise *τ_I_* is objectively low.

While these general trends hold across different values of the correlation *ρ_I_* (see panels D and E of Figure 3) we still note this parameter’s importance. In general, *ρ_I_* shapes both the additional information afforded by combining *X_I_* and *Y_I_* as well as the confidence rating process itself. Briefly, one way *ρ_I_* impacts the information seeking is by increasing the step-like nature of the high *τ_I_* curves which is visible in the difference between the *ρ_I_* = 0.2 and *ρ_I_* = 0.5 settings we depict. Also somewhat visible in our figures is the fact that with lower rater noise *τ_I_* and with low accuracy, information seeking will in fact slightly decrease. Both these aspects arise from some intricacies in the way signal and noise trade off in bivariate normal distributions. Because we focus on the cognitive rather than specifically mathematical implications of our models here, we save the discussion of these aspects for the appendix (section B).

It is worth noting that all the discussed relationships between average initial accuracy and average search also hold for the relationship between average confidence and average search. Consequently, if experimenters were to observe different agents with similar average confidence but differing underlying confidence architectures, they could expect wildly different levels of average information search.

### Cue reliability and information seeking

The decision to seek out additional information should naturally not only be influenced by the quality of the stimuli we have encountered, but also by the quality of the stimuli that we will encounter in the future. With regard to the latter, there is room between two extremes: The second piece of information might always perfectly disambiguate the judgment (small *σ_F_*) or it might carry almost no information whatsoever (large *σ_F_*). While an agent will likely want to almost always consult the former, it won’t profit much from the latter.

#### Postdecisional and first-order model

The first-order and postdecisional model capture this intuition, as evident in the first-order model depicted in Figure 4A. There, we show the average seeking for different levels of accuracy afforded by the actor’s initial and final stimulus *ϕ*(*σ_I_*) and *ϕ*(*σ_F_*) while again keeping cost constant at *r_S_* = −.1. As before the agent will seek more as the initial cue becomes noisier (right to left). In turn, decreasing final cue noise (higher values of *ϕ*(*σ_F_*); top to bottom) increases the usefulness of the additional cue and with it the average information seeking for a given level of initial actor noise *σ_I_*.

**Figure 4.**
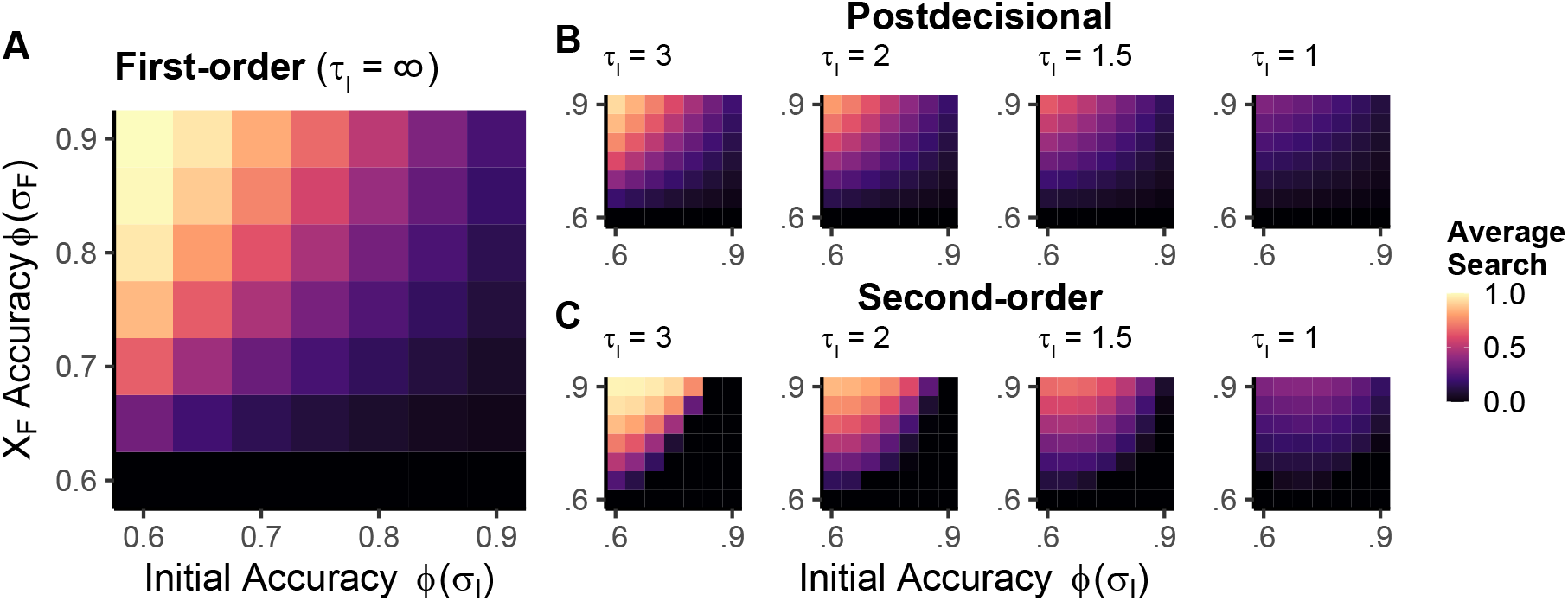
Information search as a function of cue reliability. Across models, average information seeking increases with less accurate initial decisions (ϕ(σ_I_)) and more precise final stimuli. To visualize the latter, we plot the average accuracy ϕ(σ_F_) that would be expected when encountering X_F_ in isolation. (A) This relationship is most clearly evident in the first-order model and persists in the postdecisional model (B). However, in the latter, more precise postdecisional stimuli (lower τ_I_) significantly decrease the (maximum) propensity to seek. (C) A similar pattern to the postdecisional model is visible in the second-order model, where lower values of rater noise τ_I_ also constrain the maximum average search. The transition from high to low seeking is more abrupt in the second-order model for high values of rater noise as a result of the rater’s general trust in the actor’s choice (compare especially the plots with τ_I_ = 3 in B and C). Note that the plots in Figure 3 are one-dimensional slices through the two-dimensional figures presented here. The abrupt declines in the panels in C are equivalent to that shown in panel E of Figure 3. Plots show a constant cost-level of r_S_ = −0.1, and ρ_I_ = 0.5.

Figure 4B show different levels of postdecisional noise. This produces similar patterns to the first-order model, albeit with some added complexity. The imprecision *τ_I_* of the postdecisional cue again has considerable influence on the maximum possible average seeking behaviour of the postdecisional agent. When the rater’s cue contains little noise (low *τ_I_*), almost no search is necessary. This is regardless of initial and final stimulus reliability. In turn, the information-seeking profile begins to again resemble that of a first-order agent as *τ_I_* becomes larger.

#### Second-order model

While second-order search broadly traces the postdecisional pattern arising from the interplay of the three noise parameters *σ_I_*, *τ_I_* and *σ_F_*, we can observe some further intricacies. Specifically, the second-order model’s seeking does not progress so smoothly from high to low information seeking with lower *σ_I_* and higher *σ_F_*, especially with high levels of rater noise, *τ_I_* (as we have previously observed when only varying accuracy). Rather, it begins to resemble more of a step-function as the rater knows less and less. For example, compare the highest levels of *τ_I_* = 3 in panels B and C of Figure 4. Whereas the postdecisional model smoothly transitions from high to low search, the second-order model remains with a high propensity to search relatively long before terminating search more abruptly. The reason for this can again be found in the limited information of the rater: When the rater knows little and the actor surpasses a specific relative uncertainty, the decision to sample becomes more binary across the objective accuracy range.

### Intermediate summary: Accuracy and search

In the two preceding sections, we demonstrated how search is governed by the information available to the seeker and the information expected to be gained through search. Broadly, the less information the seeker has and the more it can expect to gain from the final cue, the more it decides to seek. We highlighted how this relationship is complicated in a second-order architecture. There, the seeker doesn’t have full access to what the actor already knows. When the accuracies of the seeker/rater and the actor are particularly imbalanced, this can give rise to what looks close to step-functions in the average search profiles. In other words, the seeker either fully trusts or distrusts the actor, leading it to seek information almost always or almost never.

#### Search threshold in confidence space

Apart from the average seeking propensity, another important feature of an agent’s behaviour in our task is its internal confidence threshold for search. Put differently, how confident does an agent need to be to decide it has seen enough information? Our models allow us to investigate this phenomenon by finding the value of the rater’s internal variable for which the *Q*-values for seeking and not-seeking intersect and computing the confidence at this point. For a better intuition, compare Figure 2F, where the difference between the two values are plotted: The threshold is the point where this difference is 0. Importantly, turning this threshold into a marginalized prediction about how often an agent seeks information is not completely straightforward, as will be apparent when we later consider the underlying confidence distribution in more detail.

##### Postdecisional and first-order model

In the postdecisional model, this threshold is largely independent of the initial stimulus statistics model. Figure 5A demonstrates this by showing the confidence at which an agent would start seeking across a range of objective accuracies for the postdecisional case, for a constant final stimulus noise *σ_F_*. This confidence varies neither as a function of accuracy nor of postdecisional noise (which we do not display here).

**Figure 5.**
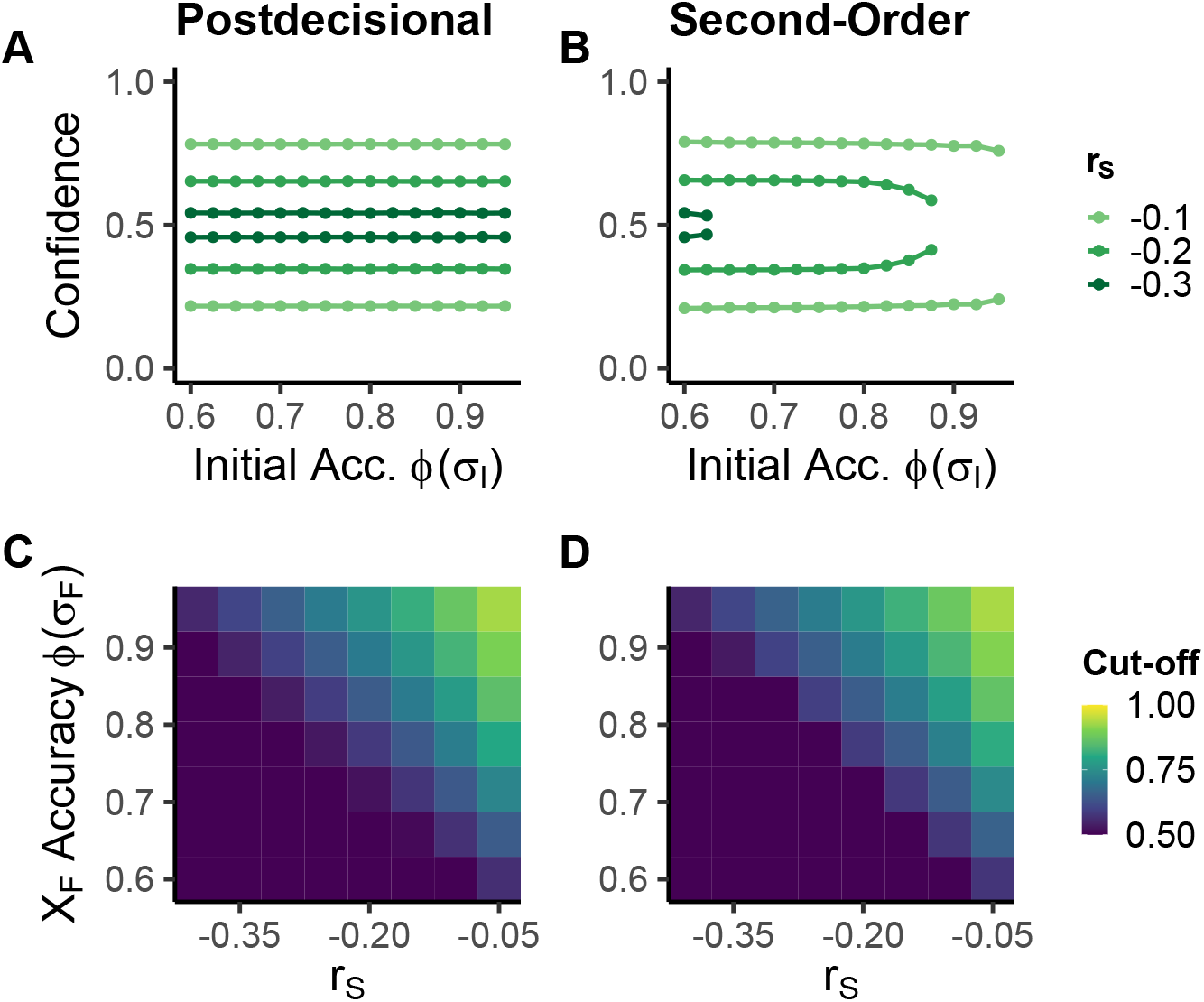
Confidence-seeking thresholds. (A,B) The confidence at which an agent stops sampling (confidence threshold) is largely independent of initial accuracy in the postdecisional and second-order model. Rather, one governing factor is the cost of the additional stimulus r_S_. By decreasing the Q–value of seeking (compare Figure 2E and F), higher costs reduce the space of confidence where it is worth probing. In the second-order model, we can also see the effect of the transition from seeking into no-seeking where the two confidence thresholds begin moving together. We set σ_F_ = 1 (ϕ(σ_F_) = 0.84), and for the second order model τ_I_ = σ_I_ as well as ρ = .5. (C,D) The two main factors governing the confidence threshold are r_S_ and the noisiness of the final stimulus σ_F_, as is visible when we plot the upper confidence threshold as a function of the two (C: postdecisional; D: second-order). Specifically, the more expensive and the less reliable the information becomes, the lower the threshold is set and the less information is sought given the same initial stimulus statistics. When the agent does not seek at all we mark the threshold as 50%. Panels C and D use σ_I_ = 3 and τ_I_ = ∞ for the postdecisional model and σ_I_ = τ_I_ = 3 and ρ_I_ = 0.5 for the second-order model respectively.

This counterintuitive result arises from the Markovian property of the first-order and postdecisional models where the confidence *c_I_* is equivalent to a belief state summarising all the previous information. In the *Q*-value computations, only this belief matters, and not how it came about. Put differently, it is unimportant whether *c_I_* was based on a large *Z_I_* and large *ζ_I_* or smaller *Z_I_* and smaller *ζ_I_*.

Rather than the initial stimulus statistics, the determining factors for the threshold are the cost of the additional information and its precision. For intuition, consider the *Q*-value functions in Figure 2E (and Figure 2F). There, the cost alters the intersection between the two *Q*-values, with higher cost reducing the space of *Z_I_* in which seeking is worthwhile and thus lowering the threshold. This influence is apparent in panel A of Figure 5. In turn, smaller *σ_F_* afford less noisy predictions of *X_F_* in Figure 2D for a given *Z_I_*, especially when this *Z_I_* is relatively unambiguous. Consequently, the *Q*(*s_I_* = 1) curve becomes steeper which leads to the intersection appearing for lower confidences. The joint influence of the final cue’s noise and cost are plotted in Figure 5C. The less expensive and the more precise the final stimulus is, the higher the boundary.

A subtle difference regarding the lower threshold appears between the first-order instance of the postdecisional model and regular postdecisional models with *τ_I_* < ∞. In the first-order version, the minimum confidence is bounded at 50% because the rater has exactly the same information as the actor. The rater will thus always endorse its decision. As a result, we only observe one “moving” boundary in the first-order model (the upper one). In contrast, when the postdecisional cue contains information, the lower bound is simply the opposite of the upper bound. This is because the net uncertainty of an initial decision made with 45% confidence is essentially the same as one made with 55%. In turn, if the rater has high confidence that the actor has made a mistake, then it can safely turn down the opportunity to acquire additional information: The actor can change its choice *a_F_* without any additional external information.

##### Second-order model

Similarly to the postdecisional model, the second-order model’s thresholds on confidence remain mostly unimpacted by the initial stimulus statistics, as is visible in Figure 5 B. There, we show the seeking threshold for a model whose rater noise *τ_I_* always equals its actor noise *σ_I_* across a range of initial accuracies. Rather, it is again the cost and noise associated with the additional stimulus that determine the threshold (see Figure 5B and D). In fact, given their differences in knowledge, it is striking that this threshold is largely equivalent between the postdecisional and second-order models, at least for low initial accuracy values. Because the second-order model also produces confidence levels below 50%, it possesses a lower threshold that mirrors the upper one, just like the postdecisional model.

As discussed above, the second-order model differentiates itself from the postdecisional model by producing behaviour where it does not seek at all. This allows us to investigate what happens in the transition to this state of uniform non-seeking. In these cases, as we can observe that with higher cost levels in Figure 5B, the two confidence cut-offs begin moving closer together until they end up meeting at 50 %. At this point, seeking stops. While the baseline threshold for low initial accuracy is thus unaffected by the initial stimulus set-up, different *σ_I_*’s can produce different initial accuracies at which seeking becomes too costly. This thus affects when the two thresholds begin moving toward each other.

The influence of the final cue noise *σ_F_* and the seeking cost *rs* on the confidence cut-off is equivalent between the postdecisional and the second-order model despite their somewhat different *Q*-value computations (compare Figure 5C and D). This is because the second-order’s additional task of predicting the *X_I_* value is required to evaluate the *Q*-values for and against seeking. The net effect is that this extra step does not impact the threshold. The constant threshold again holds for most of the range of *σ_I_* and *τ_I_*. However, we note that that it can be impacted by these parameters when nearing the second-order model’s total reluctance to seek as we discussed above.

#### Metacognitive accuracy and information search

Metacognitive accuracy broadly describes an agent’s ability to discriminate its mistakes from its successes. In our task, this manifests in distinct confidence distributions for correct and incorrect choices: Agents with high metacognitive accuracy tend to have high confidence ratings when they are correct and low confidence ratings when they made a mistake.

We can delineate two measures of metacognitive accuracy: metacognitive sensitivity and metacognitive efficiency (Fleming & Lau, 2014). Metacognitive *sensitivity* describes the aforementioned separation of confidence distributions, with less overlap between the two functions a hallmark of high metacognitive sensitivity. In our framework, this sensitivity is largely governed by the quality of the rater’s information, *τ_I_*, with higher values of *τ_I_* resulting in lower metacognitive sensitivity.

While metacognitive sensitivity provides a useful marker of the quality of an agent’s metacognition, it is often confounded with objective accuracy. Easier tasks allow more insight into the quality of our decisions – such that when objective (e.g. perceptual sensitivity) is high, metacognitive sensitivity also tends to be high (Fleming & Lau, 2014). Metacognitive *efficiency* controls for this link between objective and metacognitive sensitivity by normalizing the latter by the former. This statistic is expressed as a ratio, with values less than 1 indicating metacognitive hyposensitivity, where metacognitive sensitivity is worse than would be expected based on objective performance, and values greater than 1 indicating metacognitive hypersensitivtiy, in which case metacognitive efficiency is higher than expected based on objective performance (Fleming & Daw, 2017; Fleming & Lau, 2014).

The fact that the rater has different, possibly additional, sources of information from the actor is what licenses varying metacognitive efficiencies. The different models operationalize this slightly differently. In the postdecisional model, metacognitive efficiency can be expressed through the ratio *σ_I_*/*ζ_I_*. The larger this ratio, the more additional information the postdecisional rater has, and the higher its metacognitive efficiency. The metacognitive efficiency of the second-order model is determined by *σ_I_/τ_I_* (for a constant *ρ_I_*), again because of the restricted informational access of the second-order model. ^4^

##### First-order and postdecisional models

To understand better the relationship between seeking and metacognitive accuracy, we first need to recapitulate in detail how metacognitive accuracy arises in our models. To illustrate this better, we plot distributions of confidence ratings conditioned on accuracy in Figure 6 B, C, D and E. These illustrate the overlap between the distributions of confidence ratings for correct and incorrect answers.

**Figure 6.**
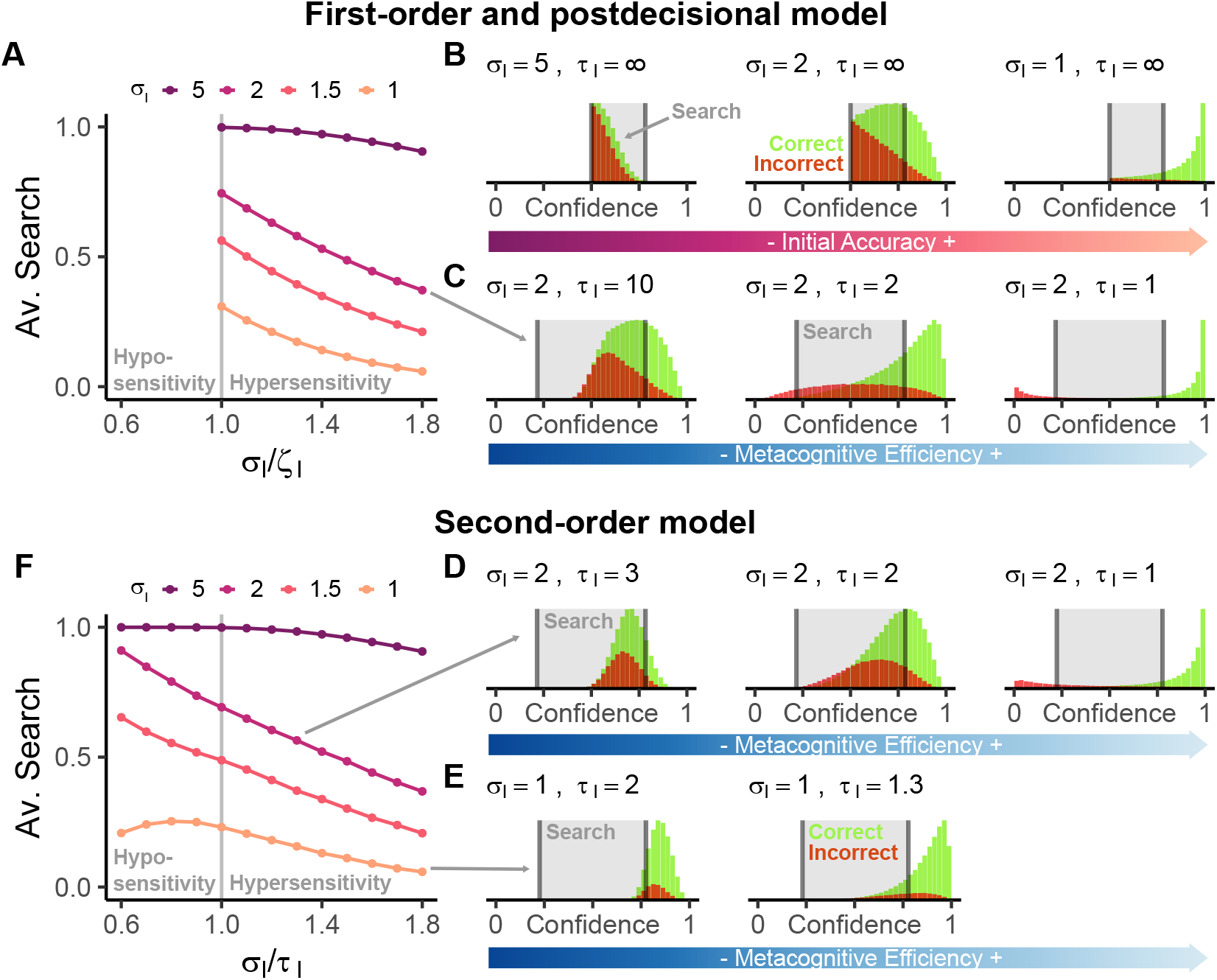
Metacognitive sensitivity, efficiency and search (A,D) Seeking as a function of the metacognitive sensitivity. (A) As the metacognitive efficiency in the postdecisional model (σ_I_/ζ_I_) increases, the need to seek more information decreases. (Note that metacognitive hyposensitivity is not possible within the postdecisional model, in which either the same or additional information is available for the confidence rating as for the decision). (B,C,D,E) Distributions of confidence ratings after correct and incorrect decisions with areas in which an agent will seek displayed in grey. (B) In the first-order model (or a postdecisional model with τ_I_ = ∞), metacognitive sensitivity is tied to the objective accuracy (σ_I_) and thus has no independent influence on search. Confidence also has a lower bound at 50%. (C,D,E) When keeping the objective accuracy (σ_I_)fixed, noise τ_I_ associated with the rater’s additional stimulus can produce diverse confidence distributions in the postdecisional model (C) and second-order model (E,F). Notice in particular how these models can correct their own mistakes without the need for additional search when confidence is below 0.5. In contrast to the postdecisional model, the second-order model can produce metacognitive hypo-sensitivity when σ_I_ /τ_I_ < 1. (E) When initial accuracy is high and metacognition particularly inefficient in the second-order model, the confidence distribution shifts almost entirely out of the seeking-zone. (F) As a result, metacognitive hypo-sensitivity can prescribe both increasing and decreasing information search in the second-order model. Metacognitive hypersensitivity is still related to reduced search. For seeking averages and thresholds all plots use σ_F_ = 1 (ϕ(σ_F_) = 0.84) and r_S_ = −0.1. In the second-order case, ρ_I_ = 0.5.

In the first order model (panels B; *τ_I_* = ∞), objective accuracy and metacognitive sensitivity are welded together. That is, higher objective performance (lower *σ_I_*) results in more clearly distinguishable confidence distributions and thus increasing metacognitive sensitivity. By design, the ratio *σ_I_/ζ_I_* is also always 1 in the first-order model, pinning down metacognitive efficiency.

In Figure 6B, we demonstrate the relationship between metacognitive accuracy and search in the first-order model. We plot the seeking-thresholds we introduced above in black and the zone of confidence values where the agent seeks in grey. Recall that these are not influenced by the statistics of the first decision. Because the confidence distributions shift together for decreasing accuracy, this results in more search. In essence, this relationship simply recapitulates what we have seen in the first section on objective accuracy and average search. Notably sensitivity will appear to be related to decreased information search in the first-order model, but this is fully explained by the coupling of metacognitive and objective accuracy. Finally, there is no relation between search and metacognitive efficiency, as the latter is invariant in the first-order model.

In contrast to the first-order model, the postdecisional model with *τ_I_* < ∞ can produce different levels of metacognitive efficiency. Figure 6 C demonstrates this by keeping the objective accuracy (*σ_I_*) constant, but varying the quality of the rater’s information through *τ_I_*. In essence, these plots take the first order model of a given objective accuracy (*τ_I_* = 2; middle of right top row of panel B), but give the rater additional information. The impact of this additional information is clearly visible: A rater with a low *τ_I_* and thus highly accurate postdecisional information is almost perfectly able to distinguish its correct from incorrect decisions, as expressed through its confidence. The confidence for correct decisions will be very high on average whereas the confidence for incorrect decisions will almost always indicate an error, that is be below 0.5. As *τ_I_* increases, the rater’s additional information decreases, resulting in a confidence distribution very similar to the first-order model when noise is very high (leftmost plot of panel C with *τ_I_* = 10), albeit one which still allows for some confidence values that are less than 0.5.

The distributions in Figure 6C also provide insight into the relationship between metacognitive accuracy and seeking in the postdecisional model. The highly separated distributions that result from low *τ_I_*’s mean confidence is pushed outside of the thresholds on both ends of the confidence range. Mistakes will likely be accompanied by a strong error signal (very low confidence) that enables a change of mind without the need for additional information seeking. In turn, correct decisions will likely trigger confidences so high that no information seeking is deemed necessary either.

We can further investigate the relationship between objective accuracy, metacognitive efficiency and search by quantifying metacognitive efficiency in the postdecisional model as the ratio *σ_I_/ζ_I_*. Importantly, the ratio is always equal to or greater than 1, because the postdecisional stimulus set-up only allows for additional knowledge (metacognitive hyper-sensitivity) but not for reduced knowledge (metacognitive hyposensitivity). Figure 6 A illustrates the effect of this hypersensitivity on search. We again observe a main effect of decision accuracy as governed by *σ_I_* (differently shaded lines), but can now see the additional effect of metacognitive efficiency: Higher levels of metacognitive efficiency give rise to reduced search on average.

##### Second-order model

The rather diverse confidence distributions produced by the second-order model are visible in panels E-F of Figure 6. In E, we again hold the objective accuracy at *σ_I_* = 2, but vary *τ_I_*. The rater noise importantly plays a different role in the second-order model, because the rater does not have direct access to the actor’s cue *X_I_*. We again see how low values of *τ_I_* give rise to clearly distinct confidence distributions and increase the chance of successful error monitoring. However, in addition the second-order model also allows for metacognitive hypo-sensitivity, when *τ_I_* > *σ_I_*. In these cases, the rater has less information than the actor. The consequences of this are visible when comparing the first-order plot with *σ_I_* = 2 plots in panel B to the second-order plot with *σ_I_* = 2 and *τ_I_* = 3 in panel D: The second order model’s two distributions are less distinguishable than the first-order model because of the hyposensitivity produced by *τ_I_* > *σ_I_*.

In general, the relationship between metacognitive accuracy and seeking holds in the second-order model. The increased levels of metacognitive insight resulting from lower values of *τ_I_* push the confidence distributions out of the zone defined by the aforementioned seeking threshold. This results in lowered search. Consequently, the effects of metacognitive hypersensitivity on seeking in the second-order model are comparable to those in the postdecisional model (see Figure 6F) – when keeping objective accuracy constant, higher efficiency again results in less need for additional information. The trend continues into metacognitive hyposensitivity. However, striking non-linear effects appear. These are again triggered by the specific knowledge states of the second-order actor and rater: Recall that if the actor is very accurate, and the rater has less knowledge, the agent’s confidence will begin to be relatively constant. In essence, the rater will begin to always trust the actor. This is visible in the leftmost plot in panel F, where *σ_I_* = 1 and *τ_I_* = 2. Here, the rater will know significantly less than the actor. The confidence ratings will thus closely congregate around the actor’s average accuracy, *ϕ*(*σ_I_*) = .84, which represents the rater’s best guess given its limited knowledge. This in turn will lead to most of the confidence ratings to be above the confidence threshold and reduce the average information seeking in comparison to a rater with more information. This effect of higher metacognitive efficiency is particularly visible in the panel E of figure 6: With the higher metacognitive efficiency of *τ_I_* = 1.3 compared to *τ_I_* = 2, there is more confidence mass within the seeking interval.

In summary, this means that search can be reduced in the second order model through two distinct mechanisms. When a second-order agent becomes more metacognitively hypersensitive, it will seek less because it has more information. However, counterintuitively, a second-order agent might also seek less when it is metacognitively hyposensitive, but this time because it has *less*, or to be more precise too little, information.

#### Intermediate summary: Confidence and search

In the preceding two sections, we discussed the intricate relationships between confidence and search. In both models, the threshold of confidence at which search is triggered is largely independent of the initial stimulus characteristics due to its (quasi-)Markovian property. Rather, the zone in confidence space where seeking is adequate is governed by the cost and precision of the additional information that can be sought out.

The confidence thresholds are however crucial when considering the confidence distributions that fall within, or outside them. In both the postdecisional and the second-order models, metacognitive hypersensivity shifts confidence outside of the seeking zone, reducing search. In the second-order model, hypo-sensitivity can trigger both increased and decreased search by either shifting more confidence into, or above the seeking zone.

#### Final accuracy

The accuracy of its ultimate, overall judgement, *a_F_*, constitutes a last crucial aspect of of an agent’s behaviour in our task. This final accuracy of course depends on the decision of the seeker – but in a potentially complex manner, because the seeker’s decisionmaking in turn is partially a function of its estimates of the benefits for this final accuracy of further search. We show these relationships in Figure 7.

**Figure 7.**
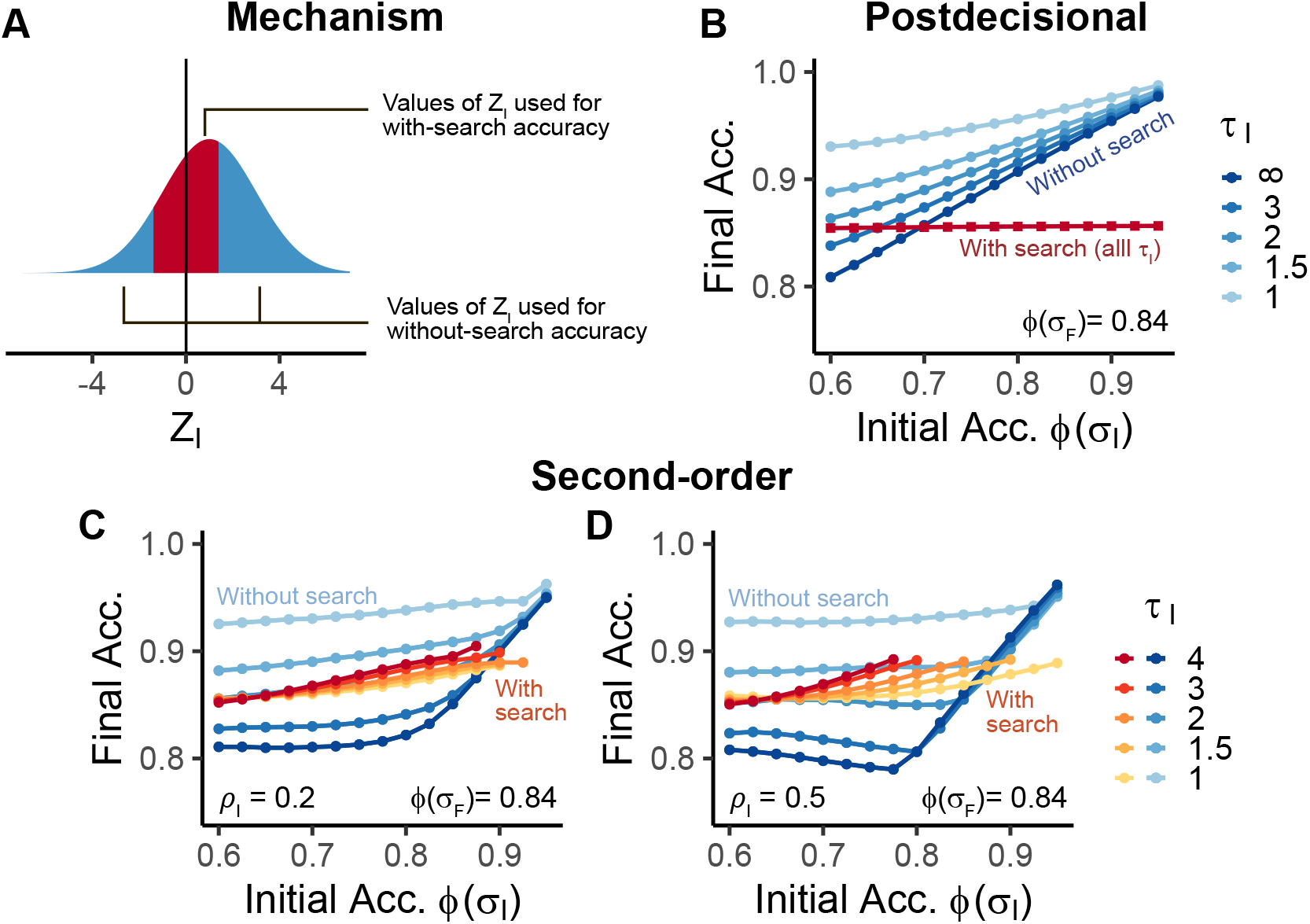
Final accuracy conditioned on search. (A) Values of Z_I_ that get passed on to the final accuracy with (red) and without seeking (blue) in the postdecisional model. When seeking, the Z_I_ values tend to be ambiguous whereas when the agent decides against search, the values tend to be more extreme and therefore offer better accuracy. (B) As a result, in the postdecisional model, the final accuracy with seeking (red) is independent of the initial stimulus. The final accuracy without seeking (blue to purple) is governed by both the accuracy afforded through the initial decision as well as the extra information contained in the postdecisional cue Y_I_. (C, D) Because the seeking decision is made without consideration of the action variable X_I_, the final accuracy differs in the second-order model. The final accuracy after seeking receives a boost through unambiguous values of X_I_ that slip through. This fact also lets the final accuracy without seeking remain relatively stable until the agent doesn’t seek any information at all, at which point it becomes a function of the initial accuracy. B - D fix final stimulus noise and cost at (ϕ(σ_F_ = 1) = 0.84)) and r_S_ = −0.1.

We distinguish between two kinds of final accuracy: The final accuracy when the agent decided not to seek additional information, and the final accuracy when the agent decided to do so. For brevity, we will in the following refer to these as the without-search accuracy and the with-search accuracy.

##### Postdecisional and first-order model

To understand how the final accuracy comes about, we need to consider what kind of cues are available to the actor at the final time point (see Figure 7A). Recall that in the postdecisional model, seeking is a function of *Z_I_*: Ambiguous values of *Z_I_* (e.g. *Z_I_* = 0) will give rise to seeking, whereas extreme values of *Z_I_* (e.g. *Z_I_* = 3) will already come with enough confidence to make information search unnecessary. Thus, as seen in Figure 7A, with-search final decisions will be made with ambiguous, more intermediate, values of *Z_I_*. In contrast, without-search final decisions will only be based on more extreme values.

This division of the *Z_I_* clearly impacts the final accuracy: In the without-search case, accuracy will be higher than would be expected when making a decision on the entire *Z_I_* space (*ϕ*(*ζ_I_*)). This increase is triggered because the error-prone, intermediate *Z_I_* values are excluded (by the fact of seeking). In contrast, in the with-search case, the actor will have relatively poor information before receiving *X_F_* due to the ambiguity associated with these intermediate values. The results of this division are visible in Figure 7C where we show the two final accuracies as a function of the initial accuracy and the rater noise *τ_I_*.

Let us first inspect the with-search accuracy, pictured in red. Strikingly, this is not influenced by either the initial accuracy or *τ_I_*. The reasons for this lie in the aforementioned stimulus set-up and relate to the confidence threshold: Because the agent only has relatively ambiguous values *Z_I_* before it receives *X_F_*, it will not have a strong preference for either option prior to search. This in turn means that the accuracy of the final stimulus *σ_F_* is the crucial determinant of the with-search accuracy. For example, in Figure 7B, *σ_F_* would afford an accuracy of around 84 % (*ϕ*(*σ_F_*) = 0.84), and the without-search accuracy is only marginally higher. This slight boost over the accuracy afforded by a solitary *X_F_* is in fact governed by the cost, which if lower, increases the range of *Z_I_*’s passed onto to seeking and therefore decreases the quality of information prior to the receipt of *X_F_*.

In contrast, the relationship between initial accuracy and final without-search accuracy (plotted in blue) is linear – at least in the first order model. That is the without-search accuracy is as accurate as the initial accuracy plus an additional boost. This again results from the *Z_I_*’s available which tend to be less ambiguous when the agent does not seek. Introducing additional information through *τ_I_* < ∞ strongly modulates the without-search accuracy. This is again the case because in these cases, the agent makes its decision not based on a stimulus with *σ_I_* but on a combined stimulus with *ζ_I_* which will always be more precise than *σ_I_*. Low noise in the postdecisional information will thus considerably boost the final accuracy through the aforementioned capability for error monitoring.

Curiously, the model sometimes produces a behaviour where it makes worse decisions with additional information than without it. At first, this can appear implausible: information should serve to increase performance. However, the agent of course has to balance the gained accuracy with its cost, triggering it to seek out information only when it is more likely to have made a poor decision.

##### Second-order model

The limited informational access of the second-order model becomes especially crucial when investigating its final accuracy. Recall that the postdecisional seeker has access to *X_I_*, and so already fully knows the quality of the final decision if it were not to seek. In contrast, the second-order seeker is less well informed. It only has access to the rater variable *Y_I_* and can make noisy inferences about the actor variable *X_I_* based on the actor’s decision. The seeker thus lacks the perfect insight afforded by the postdecisional model. As a result, seeking is only a function of *Y_I_* in the second-order model rather than the full *Z_I_* in the postdecisional model (compare Figure 1, panels D and E). As a result, the stimulus space is not, as in the postdecisional model, divided along the crucial variable for the without-search accuracy (*Z_I_*), but only along part of it, *Y_I_*.

The key problem for the second-order seeker resulting from its limited access to *X_I_* is that potentially unambiguous *Z_I_* values can slip under its radar. As an example, picture the extreme case when the rater obtains a relatively ambiguous cue (*Y_I_* = 0.3). Under most parameter combinations, this will result in low confidence and trigger the agent to seek. We can broadly think about two possible cases based on this: In one case, the actor itself might have received an ambiguous cue (e.g. *X_I_* = 0.1). In this case, in the counterfactual scenario where the agent would not have searched, its final decision would have been based on a rather ambiguous *Z_I_*. Here, seeking would have been a good decision. In another case, the actor might in fact have observed a very distinct cue (*X_I_* = 3). In this case, the actor would have already had a rather unambiguous cue *Z_I_* for a final decision in the counterfactual non-seeking scenario. Here, seeking wouldn’t be of much benefit. Whereas the postdecisional agent would know this and thus not seek, the second-order rater has no access to *X_I_* and will thus sometimes sample even though it might not have been necessary.

This divergent knowledge results in a different pattern for the with-search accuracy, with influences for both *σ_I_* and *τ_I_*. The less relative insight the seeker has (higher *τ_I_*) the more the *X_I_* “leakage”, which is visible in panel C of Figure 7. Recall there that a constant level of *τ_I_* results in decreasing metacognitive efficiency when increasing the objective accuracy. As a result, more *Z_I_*’s which would lead to no seeking on the part of a postdecisional agent with full insight are assigned to seeking by the second-order model. This “unnecessary” seeking increases the with-search accuracy until the highly metacognitively inefficient agent stops seeking entirely, as was visible in the average seeking figures (Figures 3 D & E).

The *X_I_* leakage inherent in the second-order model’s seeking computations also affects the without-search final accuracy, but to its disadvantage. Specifically, the unnecessarily good values of *X_I_* included through the myopic seeking are now no longer available to the actor when the final decision is made without search. The final accuracy thus does not increase with increasing initial accuracy while the agent still seeks (again, compare Figure 3). In fact, under certain stimulus configurations, the without-search accuracy can even slightly decrease as a result of the good *X_I_* being “stolen” by the seeking. It also worth noting that the amount of additional decision information afforded by *X_I_* also decreases with heightened correlation *ρ* reducing the additional information available.^5^

The without-search final accuracy begins to fall into a linear relationship with the initial accuracy once the agent entirely stops seeking. It is then simply a function of *ζ_I_* (*ϕ*(*ζ_I_*); by contrast with the postdecisional model where it is greater than *ϕ*(*ζ_I_*). Before then, the baseline without-search accuracy is governed by *τ_I_*, again as a result of the increasing capability for error monitoring that comes along with increasing metacognitive sensitivity. The ignition point of this increase is governed by the baseline rater noise *τ_I_* which impacts when the seeker will stop seeking entirely.

As before, the main patterns remain intact when altering *ρ_I_* in the second-order model (compare panels C and D). However, some additional subtleties arise which we will show in the appendix (section B). Briefly, it is worth noting that the *X_I_* leakage is higher for the increased *ρ_I_* because in this case more information is sought and the average seeking curve resembles more of a step-function.

### Discussion

Computational models of metacognition have recently been highly successful in explaining many intricate facets of human confidence, including error monitoring and varying degrees of metacognitive sensitivity (Fleming & Daw, 2017; Rahnev et al., 2020; Yeung & Summerfield, 2012). However, it has long been noted that metacognitive monitoring exists to guide subsequent control of behaviour (Nelson & Narens, 1990), such as knowing when to invest time and effort in studying new material or seeking new information (Desender et al., 2018; Goupil, Romand-Monnier, & Kouider, 2016; Metcalfe & Finn, 2008; L. Schulz et al., 2020). How these two processes of monitoring and control interface has attracted less attention from computational modelers. Here, we considered the rather diverse consequences that different assumptions about the informational structure underlying confidence have for optimal search. We did so by treating the process of remunerated inference and costly information acquisition in the face of uncertainty as a simple instance of a partially observable Markov decision problem (POMDP).

We extended two different model architectures suggested by Fleming and Daw (2017), exploiting the extremely simplified version of drift diffusion-like decision making discussed by Dayan and Daw (2008). In the postdecisional models, the rating process that generates confidence judgements has access to at least the information underlying the original decision whose confidence it judges, as well as in most cases additional information (Moran et al., 2015; Navajas et al., 2016; Pleskac & Busemeyer, 2010). By contrast, in the second-order model, rater and actor share only part of each other’s information, quantified by a degree of correlation (Fleming & Daw, 2017; Jang et al., 2012). In our extension, this confidence is used to determine whether the agent should, depending on the expense of doing so, collect more information before gaining reward for a final choice.

Our results highlight how only seemingly small changes in the assumed informational architecture of acting, rating and seeking can lead to starkly different profiles of optimal search. The second-order model in particular contains a number of non-trivial and often non-linear relationships between action, confidence, and optimal information search. For example, the average willingness to search as a function of objective accuracy can resemble a step-function for some parameter values in this model. In addition, because of the specific distributions of confidence associated with the second-order models, metacognitive hypo-sensitivity can give rise to both increased or decreased information search, depending on the underlying objective accuracy.

In fact, even the basic confidence judgements produced by the second-order model can have counterintuitive characteristics in certain regimes – such as that the more the rater’s private information *contradicts* the actor’s choice, the more confident the rater can be that the actor’s decision was in fact *correct*. We mainly focused on regimes in which predictions are less unusual, in keeping with the likely psychological unreality of these extremes. However, we point the interested reader to the fuller picture in appendix (section B).

Locating other monitoring accounts relative to the Fleming and Daw (2017) versions of the postdecisional and second-order models requires a careful look at their respective informational architectures. Perhaps the most significant family which is at least subtly different includes those models that assume that evidence accumulates continually (Moran et al., 2015; Pleskac & Busemeyer, 2010; Resulaj, Kiani, Wolpert, & Shadlen, 2009; van den Berg, Zylberberg, Kiani, Shadlen, & Wolpert, 2016). These relate to our models in two structurally different ways. For an example of the first, consider what Pleskac and Busemeyer (2010) call a two-stage model. Superficially, this looks like our postdecisional model: the actor makes its decision based on one source of information *X_I_*, and the rater bases its confidence on a *Z_I_* which is *X_I_* plus some additional, independent, information *Y_I_* (whose precision is usually governed by the time that passes between the action and confidence). However, in Pleskac and Busemeyer (2010), the actor uses an algorithm based on diffusion-to-bound, and so *X_I_* is perfectly predicted by *a_I_*. Consequently, whereas our *X_I_* can be accompanied by different degrees of (first-order) certainty, the accumulation bound fixes this certainty. As a result, the rater can use the actor’s decision as a sufficient statistic for the rater’s random variable, and will know (as a function of the bounds) how accurate this decision is on average. In turn, this informational set-up for the rater is an instance of what we would call a second-order model with *ρ_I_* = 0. There, the rater also only knows the average accuracy of the actor, and receives uncorrelated evidence which it combines with *a_I_* to to form its confidence. The second structural relationship is to note that the action threshold in dynamical diffusion-style models already implements an implicit case of the optional information seeking that we study explicitly – allowing more information to be collected (typically at the expense of time) given insufficient confidence.

Finally, we did not focus on the potential neural realization of the seeker, and its interaction with the likely regions involved in acting and rating (Fleming, Putten, & Daw, 2018; Shimamura & Squire, 1986; Vaccaro & Fleming, 2018) as well as the neuromodulators involved in information search and the representation of uncertainty (Hauser, Moutoussis, Purg, Dayan, & Dolan, 2018; Vellani, de Vries, Gaule, & Sharot, 2020; Yu & Dayan, 2005). It would be most interesting to probe the most obvious substrates, such as those regions involved in model-based and goal-directed control (Daw, Niv, & Dayan, 2005; Dickinson & Balleine, 2002) or state inference (Behrens et al., 2018; Schuck, Cai, Wilson, & Niv, 2016), using signatures derived from behaviour as potential correlates of neural activity.

#### Informational flow and access

Our extensions to the postdecisional and second-order model make particular choices about how information flows after the confidence rating. That is, how is the new information *X_F_*, if collected by the actor, integrated with the actor’s (*X_I_*) and rater’s (*Y_I_*) original information to make the final choice (*a_F_*)?

In our formulation of the postdecisional model, the perfectly accumulating sequential sampling renders unreasonable anything short of the full integration of the three samples (*X_I_, Y_I_, X_F_*). The optimal computations would naturally be altered if the accumulation were lossy or affected by noise, or the rater had less knowledge about the actor.

In contrast to the full access afforded by our postdecisional account, in the second-order model, the initial actor and rater are more separate. This in turn leaves various credible possibilities for their subsequent integration. We endowed the final actor with the substantial inferential ability of calculating the rater’s variable *Y_I_* from the reported confidence. However, especially if the rater is not required to report this information publicly, this may not be possible. If the information about *Y_I_* available to the final actor is less than we assume here, then the computations for search would differ, for instance limiting the benefit of low rater noise *τ_I_*.

A related question is whether and how information is further propagated in the second-order model. Here, we stopped at the final action, but the rater could, of course, also compute its confidence in this second decision. For brevity, we have not included this here, but note how an optimal rater would now need to infer both the actor’s first and second cue based on the dynamics of the first and second decision. Even in the case of no search, the rater could, in some cases, receive additional information about the actor’s first cue by observing how the actor reacts to the rater’s initial confidence (for example whether the actor changes its mind after an error signal by the rater). This raises broader issues about an internal recursive back- and forth inference between the actor and rater. Given suggestions that the two may operate below and above the threshold of awareness respectively, there may ultimately be even broader implications (Dehaene, Lau, & Kouider, 2017).

In contrast to humans, other animals’ metacognition cannot be directly assayed with confidence ratings. Experimentalists have attempted to remedy this through paradigms that indirectly probe representations of subjective correctness, such as post-decision wagering (Kepecs & Mainen, 2012), opt-out experiments (Hampton, 2001) or neural markers (Kepecs et al., 2008; Kiani & Shadlen, 2009; Nieder, Wagener, & Rinnert, 2020). Information-seeking tasks have also seen wide use (Call, 2010; Call & Carpenter, 2001). There, an animal is hypothesized to possess a form of metacognition if it seeks information in situations of uncertainty (which the experimenter controls), a behavior that already develops in human infancy (Goupil et al., 2016). However, there is ambiguity about whether confidence-related behaviours and information search in animals reflect a capacity for explicit metacognition – the ability to form a distinct representation of confidence about one’s knowledge or performance (Birch, Schnell, & Clayton, 2020; Carruthers, 2008; Kornell, 2014). To the extent that second-order architectures map onto a richer capacity for creating and using explicit confidence representations, our computational models could allow inferences about the varieties of animal metacognition when applied to the kinds of tasks used in this domain.

#### Ambiguity, computational noise, uncertainty and normativity

Following Fleming and Daw (2017), the agents in our POMDP have primary uncertainty about the stimulus on a trial, but suffer no ambiguity (or secondary uncertainty) about the inaccuracy or correlation of their sources of information. They also make their choices and confidence ratings in a computationally perfect manner, and are aware of their own metacognitive competence. They employ no form of temporal-discounting and are risk neutral. It would be formally straightforward to weaken these assumptions, various of which have been shown to influence search (Gigerenzer & Garcia-Retamero, 2017; Sadeghiyeh et al., 2020).

The case in which subjects receive information whose accuracy they are uncertain about is common in dynamic decision-making problems where, for instance, the contrast of input stimuli may change in an unsignalled manner between trials (Fleming et al., 2018; Gold & Shadlen, 2001, 2007; Kiani & Shadlen, 2009). There has been work on this in the equivalent of the first-order case. For instance, the conventional reward-rate maximizing strategy for the drift diffusion decision-making model in which evidence accumulates up to a fixed threshold changes to one in which there is what is known as an urgency function (O’Connell, Shadlen, Wong-Lin, & Kelly, 2018; Ratcliff et al., 2016) so that if the agent discovers from the length of time it is taking to reach the threshold that the information they are receiving is not very accurate, then it can make a quick, potentially inaccurate, decision, and hope that the next problem will be easier (Drugowitsch, Moreno-Bote, Churchland, Shadlen, & Pouget, 2012). It would be possible to extend our models in a similar manner, allowing separate informational accumulations over time for actor and rater; with the seeker judging when to stop and allow the actor to perform. The added complexity would be that the explicit communication between actor and rater that we allowed (with the actor’s first action *a_I_* observed by the rater; and the rater’s confidence report *c_I_* being observed by the second actor) would have to be adjusted.

Our models focused on noise coming from the signals themselves, and so we assumed an entirely noise-free decision and confidence processes. This allowed us to pinpoint the influences of our different confidence models on search. However, it is of course also limiting. Decision noise is ubiquitous in behaviour (Mueller & Weidemann, 2008; Wilson et al., 2014), and noisy computations offer a different lens for understanding metacognitive inefficiencies (Shekhar & Rahnev, 2020) and exploration (Findling, Skvortsova, Dromnelle, Palminteri, & Wyart, 2019). In our task, consider an agent which would have the opportunity to collect very accurate information, but has difficulty translating this information to good actions (for example through a low softmax-temperature or high trembling hand parameter). The introduction of such noise might also happen before the action is taken itself, for example by forgetting that degrades information over time, lossy accumulation of evidence or through noisy computations. Agents that know about their own inaccuracies should issue confidence judgements and seeking decisions that take them into account. For instance, Moutoussis, Bentall, El-Deredy, and Dayan (2011) hypothesized that people with paranoia appear to “jump to conclusions” by refusing to gather information, because decision noise renders such collection futile (although see Ermakova et al., 2018).

A more complicated problem arises if agents are confused about, or even not fully aware of, their own metacognitive skill. We assumed a form of meta-metacognitive perfection. However, this is itself questionable – and issues about how agents tune this capacity, and its psychological and neural realizations have yet to be thoroughly examined. For example, agents might exist that are metacognitively highly accurate, but might be unaware of this skill, or have low confidence in it. Conversely, individuals might posses little metacognitive skill, but could consider themselves to be great raters, in essence a meta-Dunning-Kruger effect (Kruger & Dunning, 1999). If agents do not know whether they can trust their own confidence, this naturally also has implications for our information-seeking problem, and metacognitive control more broadly.

Valence and motivational effects impact information search over and above the purely instrumental and accuracy-focused seeking we discuss. Prominently, humans are more likely to look for information that has positive valence (Gesiarz, Cahill, & Sharot, 2019; Hart et al., 2009; Jonas, Schulz-Hardt, Frey, & Thelen, 2001; Sharot & Sunstein, 2020). In turn, we tend to be reluctant to seek information that might have negative valence, but might in fact be instrumentally useful - like the results of a medical test (Gigerenzer & Garcia-Retamero, 2017; Thornton, 2008). Our models do not accommodate these aspects at the moment. However, one might combine our purely instrumental values with internal values for certain beliefs – which may or may not be in line with the accuracy goals we specify (Bromberg-Martin & Sharot, 2020).

Even when there is no valence attached to the beliefs, empirical work in paradigms close to the one we use here suggest that humans integrate cues that favour an initial judgement more than those that disconfirm it, especially when confidence is high (Rollwage et al., 2020). Such a confirmation bias can be straightforwardly modelled within our framework and might might assist in explaining behaviour (Fleming et al., 2018). Recent simulation work (Rollwage & Fleming, 2021) has shown that this an apparent confidence-induced confirmation bias can in fact be adaptive when an agent posseses second-order metacognitive hypersensitivity. Notably, Rollwage and Fleming (2021) used a different information flow for the final decision. However, this still raises interesting questions about what constitutes optimality in both the passive and active sampling of information.

Also weakening the tie to normativity are recent empirical findings that human confidence based on choices with more than two options does not necessarily resemble the full Bayesian posterior, but rather tracks the difference between the two most likely options (Li & Ma, 2020). This has interesting implications for more complex choices, and it will be important to consider how search manifests in these settings. Notably, our model already has quite a number of parameters and it is unclear whether all these cognitive processes might be distinguishable in behaviour (Wilson & Collins, 2019). Targeted experimental manipulations of these factors might thus provide better insights.

Transcranial magnetic stimulation (TMS) (Fleming et al., 2015; Rounis, Maniscalco, Rothwell, Passingham, & Lau, 2010; Shekhar & Rahnev, 2018) or pharmacological manipulations (Clos, Bunzeck, & Sommer, 2019) are able to create dissociable effects on action and confidence. Metacognition can also be trained (Carpenter et al., 2019) and there are task conditions which selectively impact action and monitoring (Bona & Silvanto, 2014; Desender et al., 2018; Graziano & Sigman, 2009; Spence, Dux, & Arnold, 2016; Vlassova, Donkin, & Pearson, 2014). Investigating how search would manifest following such manipulations might provide key insights into the interplay of metacognitive monitoring and control and their underlying computations.

Some neurological (Del Cul, Dehaene, Reyes, Bravo, & Slachevsky, 2009; Fleming, Ryu, Golfinos, & Blackmon, 2014; Goldstein et al., 2009; Persaud, McLeod, & Cowey, 2007; Shimamura & Squire, 1986) and psychological disorders (David et al., 2012; Hoven et al., 2019; Rouault, Seow, Gillan, & Fleming, 2018) as well as aging (Palmer, David, & Fleming, 2014; Weil et al., 2013) specifically affect an individual’s metacognition but leave their ”object-level” abilities relatively untouched. These would have implications for information search. For instance, agents might over- or under-estimate the usefulness of the second cue or have higher thresholds for stopping to seek, or keep on returning to check that some action (such as turning off a gas stove) has been completed (Hauser et al., 2017; Tolin et al., 2003).

Experimental manipulations of or individual differences in metacognition might provide one way to disentangle postdecisional from second-order computations in actual behaviour. However, there are other specific aspects of second-order computations that warrant further investigation. Most prominently, computing second-order confidence relies on observing the actor’s decision (Fleming & Daw, 2017) and its insight is curtailed when it can not do so, a corollary also supported by empirical evidence (Pereira et al., 2020; Siedlecka, Paulewicz, & Wierzchoń, 2016). Future experiments could follow up on this by varying whether participants make an initial decision. When participants only rate their confidence but do not perform an action, this should lead to reduced metacognitive insight, and the optimal seeking computations would be more akin to a first-order model. It would be especially interesting whether such conditions could give rise to the step-like average seeking curves when varying the underlying object-level accuracy.

#### Links to other types of information search and metacognitive control

Here we addressed a very restricted information-seeking problem. In other laboratory tasks or in real world situations, information seeking is itself often embedded in more complex decision-making tasks (Mobbs, Trimmer, Blumstein, & Dayan, 2018; E. Schulz et al., 2019). For example, in reinforcement learning problems with several options of unknown value, agents face an exploration-exploitation dilemma (E. Schulz & Gershman, 2019; Sutton & Barto, 2018). The essence of this dilemma is deciding whether to pick the option currently thought to be most valuable (exploit), or to sample from another option which might end up being better (explore). Similar to our problem, this also requires agents to balance the acquisition of information with some (opportunity) cost.

Theoretical treatments of (optimal) exploration (Gittins, 1979; Schwartenbeck et al., 2019; Sutton & Barto, 2018) and empirical investigations (Boldt et al., 2019; Speekenbrink & Konstantinidis, 2015; Wilson et al., 2014; Wu et al., 2018) of human exploration also highlight the key role of uncertainty in this decision problem. For example, the widely used Upper Confidence Bound exploration strategy (Sutton & Barto, 2018) drives agents to choose options about which they are more uncertain. These models almost always consider uncertainty in what we would characterise a first-order computation – at most wondering about the effect of different prior distributions over unknown quantities. It would be interesting to think about the equivalent of postdecisional and second-order models – where agents could gain some extra, partially independent, information about the quality of their actions, for instance by observing other agents (Zhang & Gläscher, 2020). It might then be possible to use the sort of methods we have discussed to draw out the implications for exploration.

Outside of areas related to information acquisition, confidence also plays a key role in controlling other processes. For example, cognitive offloading (Gilbert et al., 2020; Hu, Luo, & Fleming, 2019; Risko & Gilbert, 2016), such as setting reminders, is closely tied to our subjective feeling of future success. Humans also prioritize the completion of different tasks as a function of their confidence (Aguilar-Lleyda, Lemarchand, & de Gardelle, 2020) and use confidence to decide adaptively when to deploy attention (Desender, Boldt, Verguts, & Donner, 2019; van den Berg et al., 2016). On a longer time horizon, confidence also shapes learning (Bjork, Dunlosky, & Kornell, 2013; Metcalfe & Finn, 2008). Here, computational modelling has shown, that, on the one hand, we learn from our local confidence about our own broader skills (Rouault, Dayan, & Fleming, 2019). On the other, we use momentary estimates of uncertainty to steer how much we learn from errors (Behrens, Woolrich, Walton, & Rushworth, 2007; McGuire, Nassar, Gold, & Kable, 2014; Vaghi et al., 2017). Investigating these phenomena computationally through a more detailed and integrated model of metacognitive monitoring and control might provide insights into both their function and dysfunction.

Whether in our paradigm or in exploration-exploitation, the collection of information serves to increase an agent’s reward and thus has a direct instrumental purpose. However, there is also a large literature dealing with what at first glance appears to be noninstrumental information seeking. Such “curiosity” for seemingly (at least currently) reward-irrelevant information has long been a puzzle to experimentalists and theoreticians (Gottlieb & Oudeyer, 2018; Iigaya, Story, Kurth-Nelson, Dolan, & Dayan, 2016; Kidd & Hayden, 2015; Kobayashi, Ravaioli, Baranès, Woodford, & Gottlieb, 2019). As in instrumental information search, confidence often plays a key role in the treatment of such behaviour, although its role is contested. Whereas some propose a monotonic relationship between confidence and curiosity similar to our instrumental results (Berlyne, 1950; Lehman & Stanley, 2011), others argue that intermediate levels of confidence are most conducive to curiosity (Baranes, Oudeyer, & Gottlieb, 2014; Kang et al., 2009; Kidd, Piantadosi, & Aslin, 2012).^6^ Others have attempted to reconcile these two perspectives (Dubey & Griffiths, 2020). These various models might benefit from the sort of explicit treatment of the underlying confidence that we have discussed.

In the real world, information is often not solely provided by faceless sources, but by other agents with their own intentions. Over and above just being noisy (and indeed nosey), such social sources might have their own biases and interests of which successful agents need to be aware when evaluating whether they should invest in hearing their opinion and using them to inform themselves (Hütter & Ache, 2016; Pescetelli & Yeung, 2020; van der Plas, David, & Fleming, 2019). This is a particular pressing issue when faced with mis- and dis-information (Lazer et al., 2018; Pennycook & Rand, 2020). Such scenarios will require adaptive metacognitive systems to make inferences not only about themselves but also about others. Theories such as cognitive hierarchy (Camerer, Ho, & Chong, 2004) or interactive POMDPs (Gmytrasiewicz & Doshi, 2004) or Rational Speech Acts (Goodman & Stuhlmüller, 2013) could be adapted to consider hierarchies of partially self-aware agents interacting with each other.

Finally, we note that hierarchies of ever more sophisticated sub-agents that model each other inside a single decision-maker constitute a form of theory of (an internal) mind that is somewhat reminiscent of these externally-directed cognitive hierarchies (Carruthers, 2009). If the internal sub-agents enjoy their own partially individual rewards – so, for instance, the rater might have an incentive to lie about its confidence if it faces an overwhelming loss for being wrong – we can expect very rich patterns of behaviour to emerge, with agents partially fooling themselves as well as others.

## Appendix A Model details

### Postdecisional model

#### Predicting *X_F_*

To predict the location of *X_F_* for the value of seeking, the seeker combines the two possible normal distribution weighted them by the associated confidence:

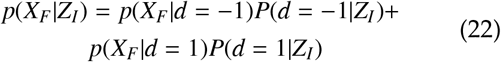

### Second-Order Model

#### Confidence

Fleming and Daw (2017) describe the computations underlying their second-order model. Here, we present them in our notation. Recall that the rater observes the actor’s decision *a_I_* and receives its own cue *Y_I_* and has to use this information to compute the probability that the actor’s decision was the correct one:

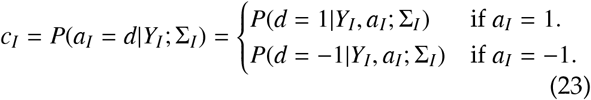

As with the postdecisional model, we apply Bayes-rule to compute this. In the following, we suppress Σ_*I*_ for clarity:

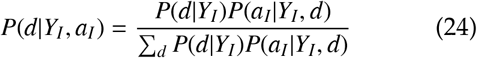

We begin teasing this apart, beginning with the second term:

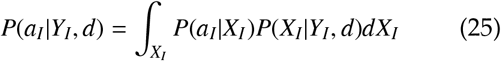

Because *P*(*a_I_*|*X_I_*) is contingent on the threshold (so that *a_I_* = 1 if *X_I_* > 0), this can also be expressed as:

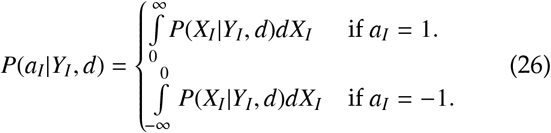

This is cumulative density function of the conditional density of a multivariate Gaussian. This conditional density of a multivariate Gaussian is itself simply a univariate Gaussian.

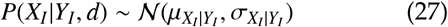

The conditional parameters of this distribution are defined as follows:

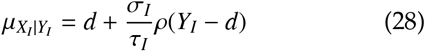

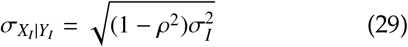

The first term is the normalized likelihood of *Y_I_* conditioned on a *d*:

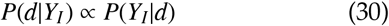

*P*(*Y_I_*|*d*) in turn equals the density of a unidimensional Gaussian with mean *d* and standard deviation *τ_I_* at *Y_I_*.

#### Optimal weighting of *X_I_* and *Y_I_* for *Y_I_* under covariance

In contrast to the postdecisonal model, we cannot simply weigh *X_I_* and *Y_I_* according to their variances when combining them to *Z_I_*. Rather, we need to take into account their covariance (Oruç, Maloney, & Landy, 2003). As a result, *X_I_* and *Y_I_* are summed with their respective weights *w_XI_* and *w_YI_*

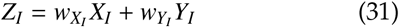

These weights are functions of the reliabilities of the cues which in turn are corrected for the correlation.

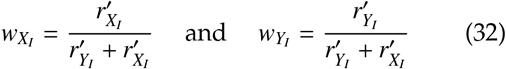

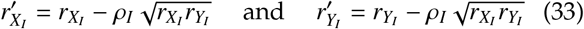

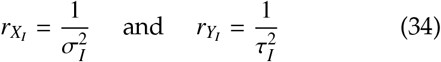

This way we can also define the standard deviation *ζ_I_* of *Z_I_*.

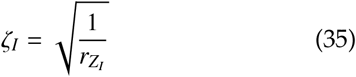

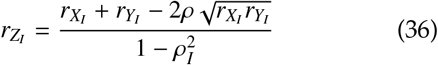

This form of cue combination can give rise to several non-intuitive results which we discuss further below.

#### Value computations

In the following, we detail the value computations in the the second-order model. First, if there is no seeking, the actor uses *Z_I_* (see above) to make its decision. The value of this combined stimulus is defined as:

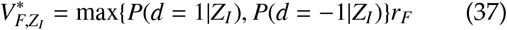

This is then used in the *Q*-value computations for the *Q*-value of not seeking (see equation 17)

However, the seeker does not know *Z_I_*, because it does not have access to *X_I_*. It therefore has to marginalize out this quantity

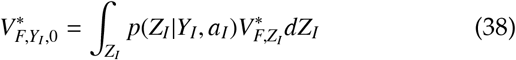

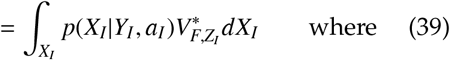

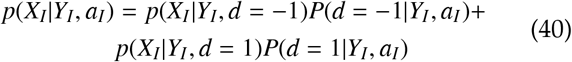

Given seeking, the actor receives *X_F_* (again as per equation 10) which it combines with *Z_I_* to form a joint variable *Z_F_* (see equation 12). This variable can then again be compared against a threshold for *a*_*F*,1_. Given this set-up, we can now consider the values that go into the individual *Q*-value computations.

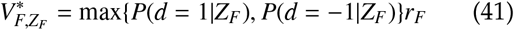

Similarly to the first-order and postdecisional models, the seeker does not know all the variables underlying *Z_F_*, when it decides whether to seek, and it also does not know *Z_I_*. Therefore, it has to marginalize over them both:

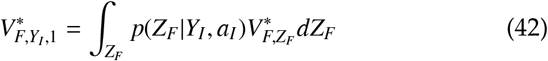

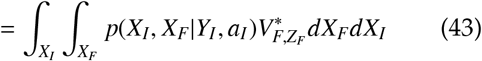

where

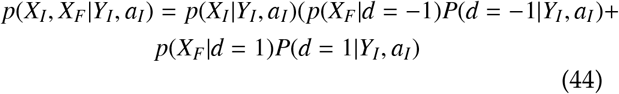

Notice how both the with- and without-search value computation contain, *P(d|Y_I_, a_I_*), or the rater’s confidence.

## Appendix B Further Second-Order Results

### Confidence and general stimulus conditions

#### Signal, noise correlation

In the second-order stimulus condition, the correlation can give rise to counterintuitive confidence curves. This is visible in Fig B1 where we plot confidence values for a positive decision (*a_I_* = 1) varying the parameters individually. We observe a few aspects already reported by Fleming and Daw (2017):

- Panel A: Increasing the accuracy of the actor (lower *σ_I_*) increases the boost that the confidence receives through the action. If the actor is very accurate, it takes a highly negative *Y_I_* to overturn the decision.
- Panel B: Higher rater noise (*τ_I_*) means the confidence curves will be less well-tuned.
- Panel C: Higher correlations (*ρ_I_*) also results in a reduced sharpness in the confidence curves.

However, what has yet to be reported is the following: Under conditions of metacognitive hyposensitivity, that is when *σ_I_* is sufficiently smaller than *τ_I_*, and when *ρ_I_* is large enough, confidence will begin rising again with seemingly contradictory *Y_I_*’s. This is particularly visible in the rightmost panel where *ρ_I_* is most pronounced, but is also visible in the most extreme cases in panels A and B. As an example, imagine the actor has received *X_I_* = 0.5 and decides *a_I_* = 1. If the rater receives *Y_I_* = −5, this would usually be a strong error signal and the confidence in the initial decision very lower than when *Y_I_* would have had more intermediate values. However, under some parameter conditions, the exact opposite is the case: There, when *Y_I_* strongly contradicts the decision sign, confidence will in fact be higher for this very low *Y_I_* than for *Y_I_* = 0.

While we note that the marginal probability of these cases is relatively low given the underlying correlation, such a pattern is striking. The reason for it lies in the the way the two possible sources occupy the *X_I_* and *Y_I_* space and create signal and noise (compare Figure 1 E). A crucial aspect of this is the line on which the posterior based on *Z_I_* (i.e. the combination of *X_I_* and *Y_I_*) is uniform, so that *P*(*d* = 1|*X_I_, Y_I_*) = *P*(*d* = −1|*X_I_, Y_I_*) = 0.5. It is this posterior that the rater only has partial information about. The equality line subdivides the space in two zones where the likelihood of *d* = 1 is larger than the likelihood of *d* = −1 (or vice versa). Given the equal prior, this line of equality in turn is defined by the points at which the two likelihoods equal each other.

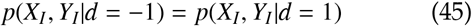

**Figure B1.**
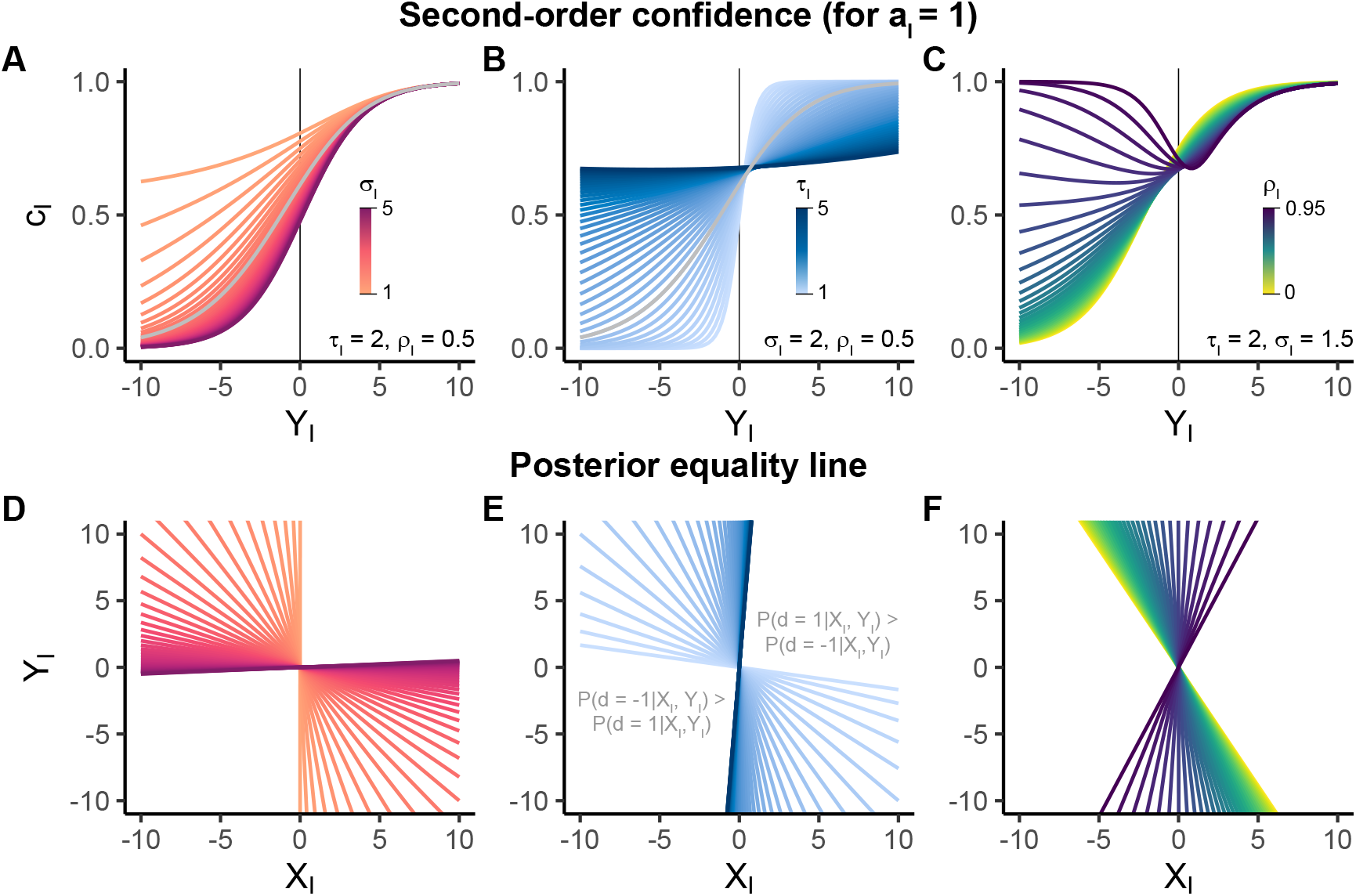
Second-order confidence across parameter regimes. (A-C) Second-order confidence as a function of Y_I_ for a_I_ = 1. In general, note how confidence for a completely ambiguous rater cue (Y_I_ = 0) doesn’t necessarily mean that confidence c_I_ will be 0.5. (A) This is mainly a function of the relationship between σ_I_ and τ_I_. (B) High values of τ_I_ for a fixed σ_I_ can lead to the confidence being less sensitive to Y_I_. When τ_I_ is particularly large in relation to σ_I_, the confidence will in fact again begin to rise for negative Y_I_’s (which intuitively contradict a_I_). Grey line in (A) and (B) highlights an equivalent parameter setting of τ_I_ = σ_I_ = 2,ρ_I_ = .5. (C) This rise of confidence with contradictory rater cues Y_I_ is particularly pronounced for high correlations ρ_I_. The rising confidence is tied to the way the correlation affects signal and noise and in extension the line on which the joint posterior P(d|X_I_, Y_I_) is equivalent between the two d. This line is plotted in (D-F). When metacognitively hyposensitive (σ_I_ < τ_I_) and when the correlation ρI between Y_I_ and X_I_ is high enough, confidence will not decrease for negative Y_I_ but rather again rise.

The two likelihoods are defined by the bivariate normal distribution’s density:

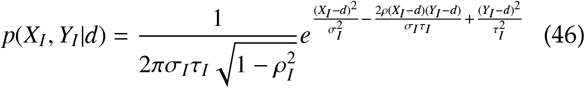

From this, we can can define the values of *Y_I_* for which the two posteriors equal each other as a function of *X_I_*:

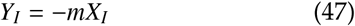

where from equation 45, we get:

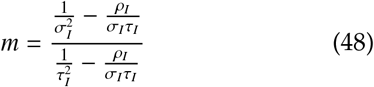

We plot this in Figure B1 D-F for a range of parameter combinations. When there is no correlation *ρ_I_* = 0, *m* (panel F) this line is defined by 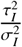 and the space is thus divided diagonally from a positive *Y_I_* to a negative *Y_I_* with the slope defined by the relationship between the two parameters. This general result holds, even when re-introducing the correlation. Importantly, what this division of space means is that for every possible actor cue *X_I_*, more positive rater cues *Y_I_* will favour *d* = 1 and more negative *Y_I_* will favour *d* = −1. Crucially however, under metacognitive hyposensitivity (*τ_I_* > *σ_I_*) this diagonal becomes steeper and steeper until it is fully vertical. This point is defined when:

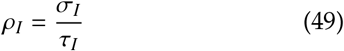

In other words, at this point, the decision rule based on *Y_I_* and *X_I_* is the same as based on *X_I_* alone – *Y_I_* thus affords no additional help with the decision. Beyond this vertical point, the space is again divided diagonally, but the dividing line now has a positive rather than negative slope. This only appears under relatively extreme parameter combinations, but will crucially flip the logic outlined above. Now, for every *X_I_*, lower values of *Y_I_* will begin providing more evidence for *d* = 1 instead of *d* = −1. This then in turn gives rise to the confidence rising with seemingly contradictory values of *Y_I_*. This phenomenon will appear once the equality lines have ‘flipped’, as is visible when comparing the confidence curves and slopes depicted in Fig. B1.

### Relationship between *ζ_I_* and *σ_I_, τ_I_, ρ_I_*

As alluded to in the main text, the joint standard deviation *ζ_I_* produced from optimally combining *σ_I_*, *τ_I_* and *ρ_I_* stands in a non-trivial relationship with its subparts.

For context, recall how the *σ_I_* and *τ_I_* are combined when there is no correlation (see equation 5). As we discussed in the main text, the maximum of *ζ_I_* is then defined by the smaller of the two standard deviations *σ_I_* and *τ_I_*. Additionally, the smaller the larger of the two is, the smaller *ζ_I_* becomes. In other words, the agent would benefit from a reduction of noise in both cases. For an illustration of this effect, see the yellow-most lines in Figure B2A that show a cue integration in accuracy space (*ϕ*(*ζ_I_*)) as a function of *ϕ*(*σ_I_*) for *ρ_I_* = 0. Notice how lower *τ_I_*’s shift the baseline upwards and how the better accuracy of afforded by *σ_I_* increases the accuracy afforded by *ζ_I_*.

In most cases of optimal cue combination, two independent sources (low *ρ_I_*) of information hold more information (lower *ζ_I_*) than two correlated sources (high *ρ_I_*). This is also the case for most parameter combinations in our scenario. Crucially however, this intuitive relationship fails for some specific combinations of values, particularly for very high correlations. This is visible in Figure B2A where for a fixed rater noise *τ_I_ lower* accuracy *σ_I_* produce *more* accurate *ζ_I_* than *higher* accuracy *σ_I_* (especially *τ_I_* = 2 in panel A and *ρ_I_* = 0.8 in panel B).

Figure B2 shows these non-monotonic relationships for a range of parameter combinations. This broadly highlights that, if parameter combinations are extreme, then there is no monotonic relationship between the three initial source parameters and the accuracy afforded by their combination (*ϕ*(*ζ_I_*).

These pattern again partially stem from how the space is optimally divided by the two sources. Specifically, when the equality line ’flips’, the posteriors get compressed differently between the two sources, allowing a better inference than in the classical separation of *X_I_*, *Y_I_* space.

The effects of this “flip” are formally analogous to the way in population codes that correlations between the activities of units can either help or hurt discrimination and decoding depending on their alignment relative to the way that signals are coded (the mean difference) (Abbott & Dayan, 1999).

### Seeking and final accuracy for the high *ρ_I_*

The two aforementioned particularities of the second-order model also impact the agent’s search behaviour and final accuracy, which we depict in Figure B3.

The fact that confidence rises again with contradictory values of *Y_I_* will result in U-shaped seeking curves for most *τ_I_*. This is because the rising confidence will favour not seeking, rather than seeking once the actor accuracy is below a specific value while keeping *τ_I_* fixed.

With regards to the final accuracy, the maximum attainable accuracy from combining *X_I_* and *Y_I_* (and *X_F_*) will be impacted by the combination of *σ_I_,τ_I_* and *ρ_I_* giving rise to *ζ_I_* (discussed above). This will for example mean that more accurate (low *τ_I_*) raters can produce less accurate final judgements than noisier (high *τ_I_*) raters.

## Appendix C Methods

We implemented our models and simulations in R. Our code will be provided as online supplemental material upon publication and hosted openly on a dedicated github repository (github.com/lionschulz/).

**Figure B2.**
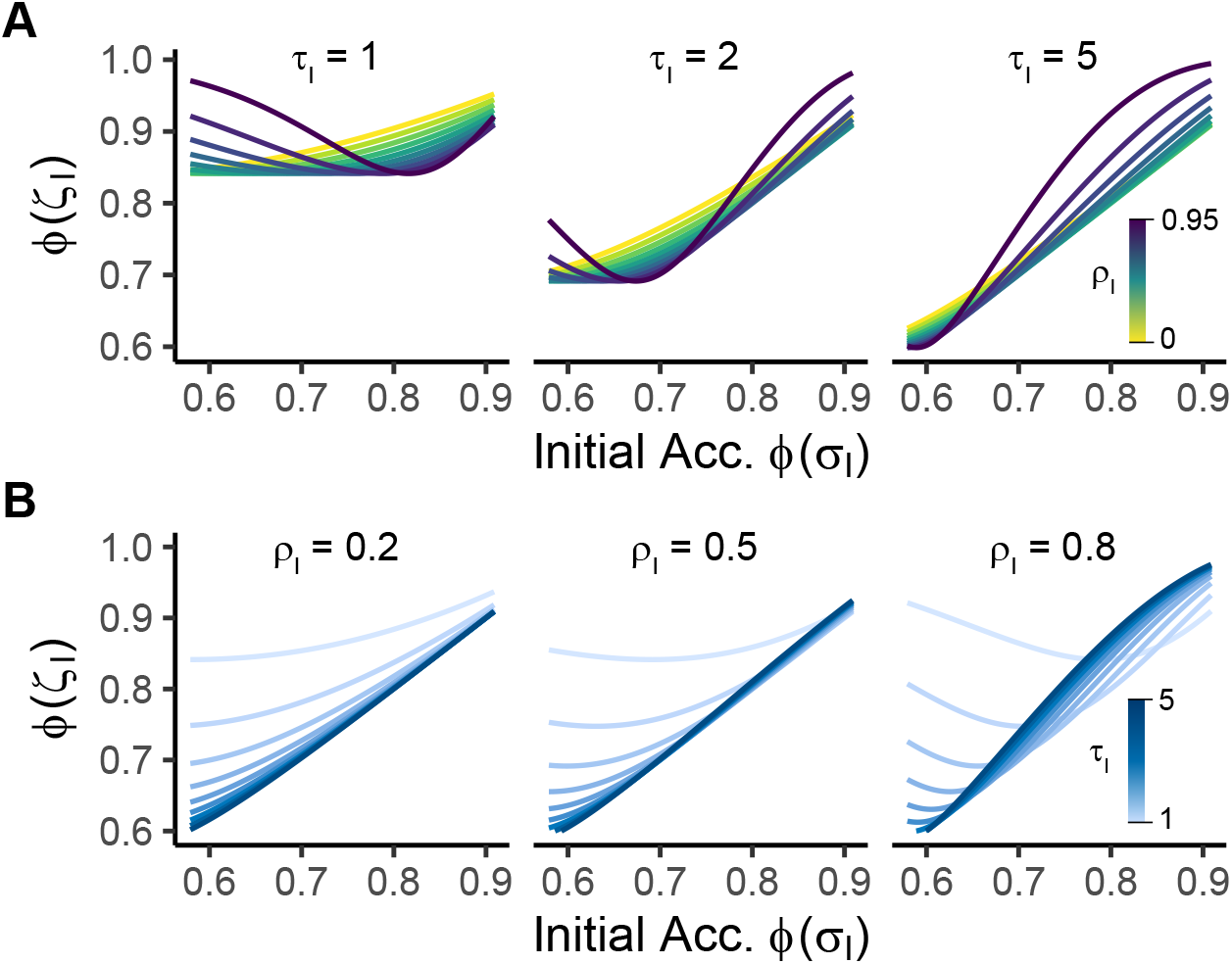
Accuracy obtainable through the standard deviation ζ_I_ of combined cue Z_I_ in the second order model. Optimally combining the parameters of the initial decision (σ, τ_z_ and ρ_I_) can give rise to non-monotonic relationships between initial accuracy and accuracy attained through ζ_I_, i.e. ϕ(ζ_I_).

**Figure B3.**
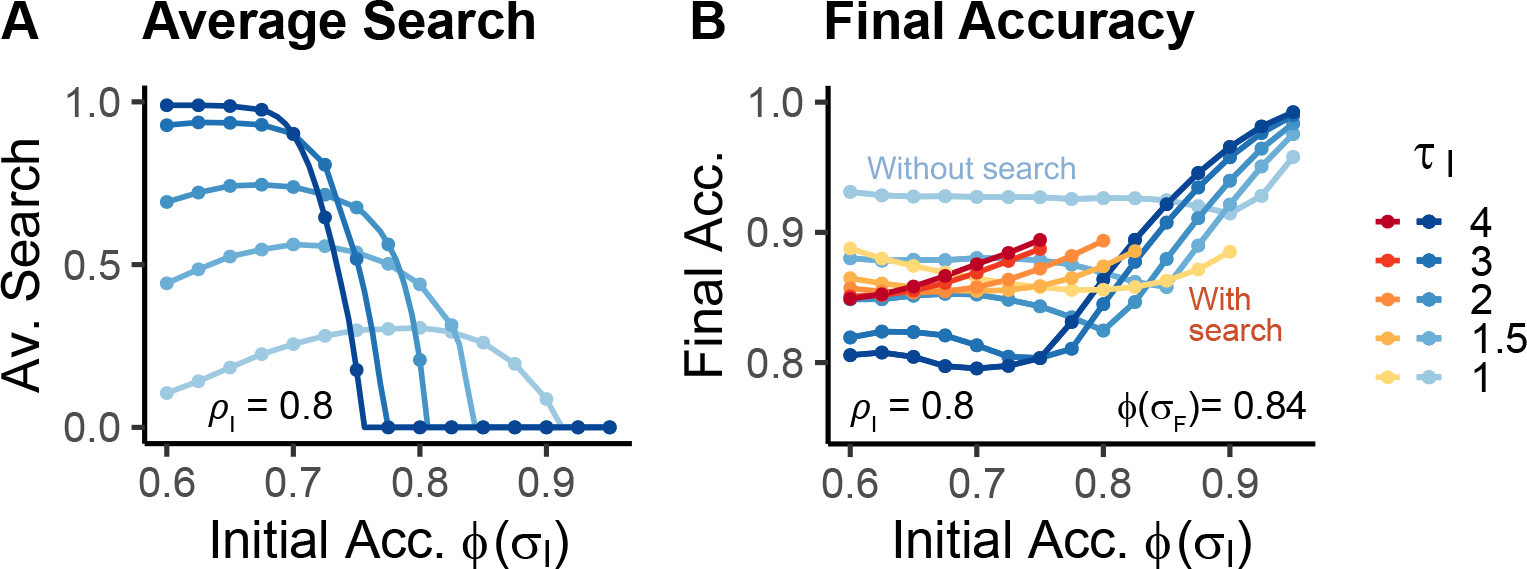
Effects of high correlation between actor and rater signal. (A) Average search by average initial accuracy and rater noise τ_I_. (B) Final accuracy by average initial accuracy and rater noise, and conditioned on whether the agent sought out information or not.

## Footnotes

1 Our postdecisional model somewhat extends Fleming and Daw (2017) but is using a formally equivalent architecture. Specifically, whereas Fleming and Daw (2017) only discuss cases where *τ_I_* = *σ_I_* (and thus 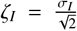), we allow the actor and noise to vary independently and describe how the two can be optimally combined. This additional flexibility enables us to subsume the first-order model within the postdecisional model and will be key in our results later when we we describe different levels of metacognitive insight.

2 We note that the specific distributions used for this simple form of second-order model have some previously unexplored peculiarities at various limiting values. Because these are not essential for our investigations, we discuss them in the appendix, section B.

3 We use this simple pay-off scheme for simplicity, but note the possibility of others (including temporal discounting). Furthermore, we assume that the agent has a linear utility function, which precludes forms of risk-aversion.

4 In the experimental literature, a plethora of measures assay metacognitive sensitivity and/or efficiency (Fleming & Lau, 2014). Most prominently, the meta-d′ statistic (Maniscalco & Lau, 2012) allows metacognitive sensitivity to be estimated within a signal detection theoretic (SDT) framework. Briefly, this approach estimates the *d′* from a first-order SDT model that best fits the observed confidence distributions. This parameter, known as meta-d′ can then be compared to the *d′* calculated from the participant’s choices to produce a ratio meta-*d′/d′*, a typical measure of metacognitive efficiency (Fleming & Lau, 2014). Both meta-d′ and meta-d′/d′ scale with our parameters *τ_I_* and, depending on the model, the ratio *σ/ζ_I_* or *σ/τ_I_* (Fleming & Daw, 2017). However, for clarity we will focus on our parameter regime in the following.

5 How *ζ_I_* stands in relationship with *ρ_I_, σ_I_* and *τ_I_* is in fact more complex under certain more extreme parameter combinations, as we discuss in further detail in the appendix B.

6 We observe inverse U-shapes under some extreme parameter settings, but stress that these are due to the signal and noise properties of the second-order model (see appendix B)

